# Fast animal pose estimation using deep neural networks

**DOI:** 10.1101/331181

**Authors:** T.D. Pereira, D. E. Aldarondo, L. Willmore, M. Kislin, S. S.-H. Wang, M. Murthy, J. W. Shaevitz

**Affiliations:** Princeton Neuroscience Institute, Princeton University; Department of Molecular Biology, Princeton University; Lewis-Sigler Institute for Integrative Genomics, Princeton University; Department of Physics, Princeton University

## Abstract

Recent work quantifying postural dynamics has attempted to define the repertoire of behaviors performed by an animal. However, a major drawback to these techniques has been their reliance on dimensionality reduction of images which destroys information about which parts of the body are used in each behavior. To address this issue, we introduce a deep learning-based method for pose estimation, LEAP (**L**EAP **E**stimates **A**nimal **P**ose). LEAP automatically predicts the positions of animal body parts using a deep convolutional neural network with as little as 10 frames of labeled data for training. This framework consists of a graphical interface for interactive labeling of body parts and software for training the network and fast prediction on new data (1 hr to train, 185 Hz predictions). We validate LEAP using videos of freely behaving fruit flies (*Drosophila melanogaster*) and track 32 distinct points on the body to fully describe the pose of the head, body, wings, and legs with an error rate of <3% of the animal’s body length. We recapitulate a number of reported findings on insect gait dynamics and show LEAP’s applicability as the first step in unsupervised behavioral classification. Finally, we extend the method to more challenging imaging situations (pairs of flies moving on a mesh-like background) and movies from freely moving mice (*Mus musculus*) where we track the full conformation of the head, body, and limbs.

## Introduction

Connecting neural activity with behavior requires methods to parse what an animal does into its constituent components (movements of its body parts), which can then be connected with the electrical activity that generates each action. This is particularly challenging for natural behavior, which is dynamic, complex, and seemingly noisy. Human classification of behavior is painstakingly slow and subject to bias – but recent methods make it feasible to automate the analysis of behavior ^1^. These include methods to track animal centroids over time ^2–4^, machine learning techniques for identifying user-defined behaviors, such as fighting or courting ^5,6^, and software to segment the acoustic signals produced by an animal ^7–9^. However, one may not know *a priori* which behaviors to analyze – this is particularly true when screening mutant animals or investigating the results of neural perturbations that can alter behavior in unexpected ways.

Recent developments in the unsupervised clustering of postural dynamics have overcome many of these challenges by analyzing the raw frames of movies in a reduced dimensional space (e.g., generated using Principal Component Analysis (PCA)). By comparing frequency spectra or fitting auto-regressive models ^10,11^, these methods both define and provide the ability to record the occurrence of tens to hundreds of unique, stereotyped behaviors in animals such as fruit flies or mice. These unsupervised methods have been used to uncover new structure in behavioral data, facilitating the investigation of temporal sequences ^12^, social interactions ^13^, the analysis of genetic mutants ^11,14^, and the results of neural perturbation ^15,16^.

While powerful, a major drawback to the aforementioned techniques is their reliance on PCA to reduce the dimensionality of the image time series. While this produces a more manageable substrate for machine learning, the modes derived from PCA come from the statistics of the images and are not related directly to any individual body part of the animal. As such, the discovered stereotyped behaviors must be labeled, classified, and compared manually through the human observation of representative movie snippets. Given the highly quantitative approach that precedes this step, it is ultimately unsatisfying and subjective for the experimenter to manually label each behavior (e.g., foreleg grooming, hindleg grooming, forward locomotion, right turns, etc.). Instead, what is desired is a mathematical representation of the relative motions of all parts of the animal that characterizes a particular behavior. Such a description would facilitate the investigation of the similarities and differences between behaviors and likely improve the behavioral identification algorithm itself.

Measuring all of the body part positions from raw images is a challenging computer vision problem. Previous attempts at automated body-part tracking in insects and mammals have relied on either physically constraining the animal and having it walk on a spherical treadmill ^17^ or linear track ^18^, applying physical markers to the animal ^17,19^, or utilizing specialized equipment such as depth cameras ^20–22^, frustrated total internal reflection imaging ^23,24^ or multiple cameras ^25^. Meanwhile, approaches designed to operate without constraining the natural space of behaviors make use of image processing techniques that are sensitive to imaging conditions and require manual correction even after full training ^26^.

To address these issues, we turned to deep learning-based methods for pose estimation that have proven successful on images of humans ^27–33^. Major breakthroughs in the field have come from adopting fully convolutional neural network architectures for efficient training and evaluation of images ^34,35^ and producing a probabilistic estimate of the position of each tracked body part ^28,30^. However, the problems of pose estimation in the typical human setting and that for laboratory animals are subtly different. Algorithms that work on human images are meant to deal with large amounts of heterogeneity in body shape, environment, and image quality, but for which there are very large labeled training sets of images available. On the contrary, behavioral laboratory experiments are often more controlled, but the imaging conditions may be highly specific to the experimental paradigm and labeled data is not readily available and must be generated for every experimental apparatus and animal type. One recent attempt to apply these techniques to images of behaving animals successfully used transfer learning, whereby networks initially trained for a more general object classification task are refined by further training with relatively few samples from animal images ^36^.

We have taken a different approach that combines a graphical user interface (GUI)-driven workflow for labeling images with a simple network architecture that is easy to train and requires fewer computations to generate predictions. Our method can automatically predict the positions of animal body parts via iterative training of deep convolutional neural networks with as little as 10 frames of labeled data for initial prediction and training. After initial *de novo* training, incrementally refined predictions can be used to guide labeling in new frames, drastically reducing the time required to label sufficient examples (∼500 frames) to achieve an accuracy of less than 3 pixels (distance from ground truth). Our framework consists of a GUI for interactive labeling of ground truth body part positions as well as software for efficient training of a convolutional neural network on a workstation with a modern GPU (<1 hour) and fast prediction on new data (up to 185 Hz). We validate the results of our method using a previously published dataset of high quality videos of freely behaving adult fruit flies (*Drosophila melanogaster* ^10^) and we recapitulate a number of reported findings on insect gait dynamics as a test of its experimental validity. We then show its applicability as a front end to an unsupervised behavioral classification algorithm and demonstrate how it can be used to describe stereotyped behaviors in terms of the dynamics of individual body parts. Finally, we show the generalizability of this method in challenging imaging conditions as well as in freely moving rodents.

## Results

Our method, which we refer to as LEAP (**L**EAP **E**stimates **A**nimal **P**ose), consists of three phases (**Fig. 1a**): *(i) Registration and alignment*, in which raw video of a behaving animal is preprocessed into egocentric coordinates; *(ii) Labeling and training*, in which the user provides ground truth labels to train the network to find body part positions in a subset of images; and *(iii) Pose estimation*, in which the network can be applied to new and unlabeled data. In the following sections, we demonstrate the power of this tool using a previously published data set of 59 male fruit flies, each recorded for one hour at 100 Hz, for a total of >21 million images ^10^. All code and utilities are available at https://github.com/leap/talmo.

**Figure 1:**
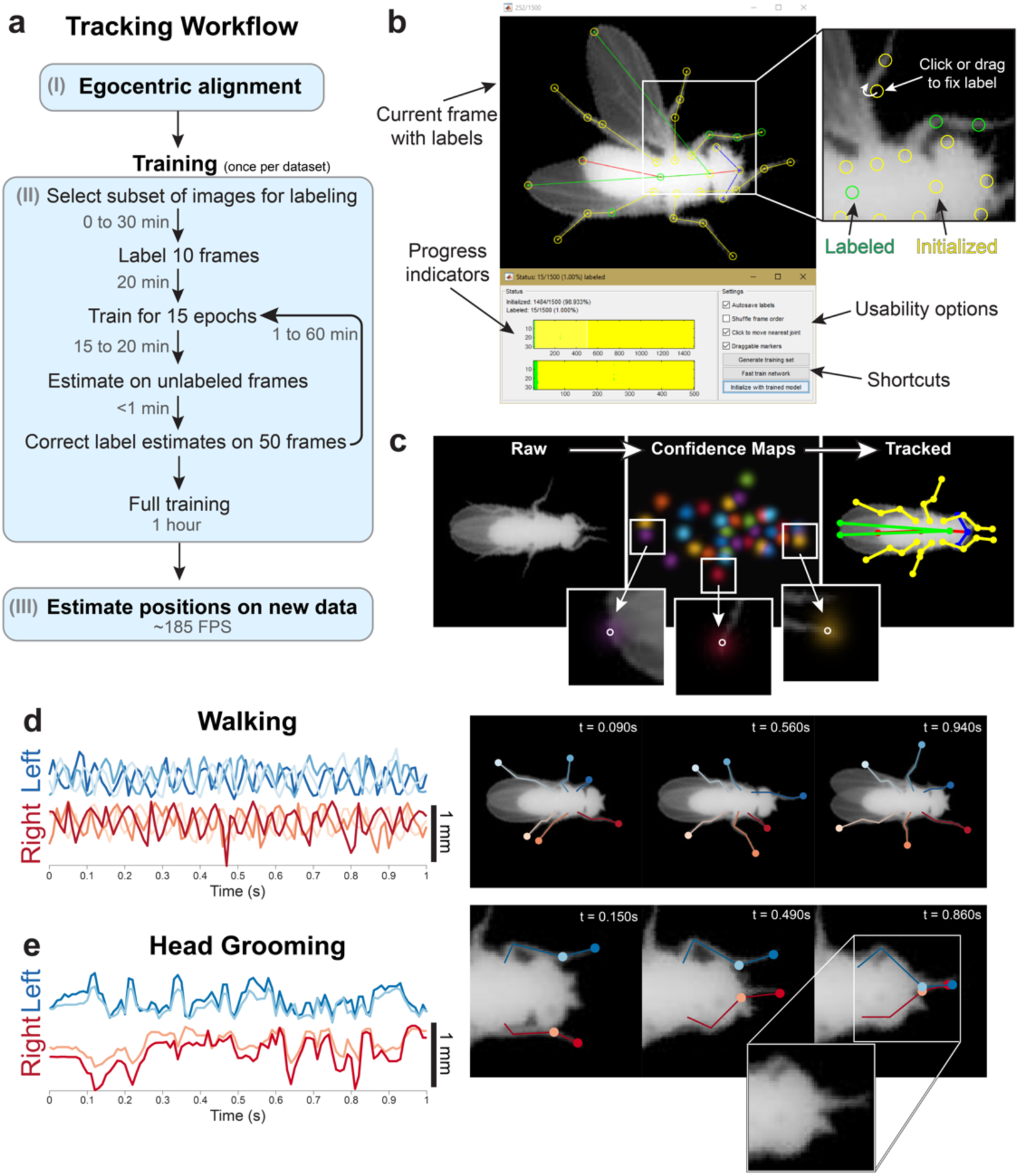
Body part tracking via LEAP, a deep learning framework for animal pose estimation. (a) Overview of the tracking workflow. In the initial preprocessing phase (I), video frames are centered relative to the animal to render the images in egocentric coordinates. In the beginning of the training phase (II), representative frames are sampled. After labeling an initial set of images, the neural network is trained and used to estimate body positions on the remaining images of the training set to facilitate subsequent correction of labels. Correcting labels takes progressively less time as the network is trained with increasingly more labeled examples. Once all training images are labeled, full training involves fine tuning the network to optimize performance. Once trained (III), estimation on new, unlabeled data is fully automated and can be performed at high speed on a GPU. (b) Graphical user interface for providing ground truth labels and correcting estimates. The software displays images in the training set with interactive markers denoting the default or best estimate for each body part (top-left). User input is provided by clicking or dragging the markers to the correct location (top-right). Colors indicate labeling progress and denote whether the marker is at the ground truth location (green) or is an estimate from network initialization (yellow). Progress indicators mark which frames and body parts have been labeled thus far, while shortcut buttons enable the user to export the labels to use a trained network to initialize unlabeled body parts with automated estimates. (c) Data flow through the LEAP pipeline. Raw images are provided as input without markers or indicators (left). For each input image, the network outputs a stack of confidence maps, a max projection through which is used here for visualization (middle). Insets overlay individual confidence maps on the image to reveal how confidence density is centered on each body part, with the peak indicated by a circle. The predicted coordinate for each body part is the peak value in each confidence map, enabling a visualization of the tracked skeleton (right). Walking behavior can be quantitatively described by leg tip trajectories. The distance of each of the 6 leg tips from its own mean position during a walking bout reveals a cyclic pattern of leg movements (left). The tracked points on the images span a diversity of poses that change over fast timescales (right). (e) Head grooming behavior can also be quantitatively described by leg tip trajectories. Position estimates are not confounded by occlusions when the legs pass under the head (right, inset).

### The Components of LEAP

#### (i) Registration and alignment

The first step in our pipeline is to extract the image region that contains the animal within the field of view of the camera, as well as its angular heading within the image. This can be accomplished using standard image processing techniques ^37,38^ or existing software packages ^2,13,39,40^. Our implementation ^10^ is provided in the accompanying code repository. This step produces egocentric, oriented bounding boxes around each fly image used to train the neural network. While this step improves pose calculation accuracy as it saves the network from being required to learn rotational invariance, we note that this can also be learned at the cost of prediction accuracy (**Supplementary Fig. 1**).

#### (ii) Labeling, training, and neural network architecture

The neural network learns to predict body part positions from a set of user-labeled images. To identify a small set of example ‘training’ images that are representative of the set of poses across the entire data set, we use a technique we refer to as *cluster sampling*. A simple random subset of the movie images are grouped via k-means clustering and then these images are sampled uniformly across groups for labeling. The grouping is based on linear correlations between pixel intensities in the images as a proxy measure for similarity in body pose. The diversity of poses represented using this method can be observed in the centroids of each of the clusters identified (**Supplementary Fig. 2**).

Poses in each training image are labeled using a custom GUI with draggable body part markers that form a skeleton (**Fig. 1b**). For the fruit fly, we track four points on each of the six legs, two points on the wing tips, three points on the thorax and abdomen, and three points on the head for a total of 32 points in every frame. These points were chosen to align with known *Drosophila* body joints (**Supplementary Fig. 3**). For every training image, the user drags each skeleton point to the appropriate body part and the program saves the label positions into a self-contained file. To enhance the size of the training image set further without the need for hand labeling more frames, we augment the dataset by applying small random rotations and body-axis reflections to generate new samples from the labeled data. As the neural network processes the raw images, the rotated and reflected images add new information that the network can use during training.

We first labeled only 10 images, and used these data to train the neural network and generate body part position estimates for the remaining images chosen via cluster sampling (see below for details on network training). When trained with only 10 images for just 15 epochs, estimation error rates were large (**Supplementary Fig. 4a-b**) but these estimates helped to decrease the time required to label each subsequent frame. We therefore repeated this procedure of alternating labeling and initializing via briefly trained network estimates at 50, 100, 250, 500 and 1000 labeled frames, decreasing the time required to label each frame from 2 minutes per frame for the first 10 frames, to 6 seconds per frame for the last 500 frames (**Supplementary Fig. 4c**). Labelling 1500 frames required a total of 7 hours of manual labeling and an additional 1.5 hours of network training (including 6 “fast” and 1 “full” training epochs).

The core component of LEAP is a deep convolutional neural network. The network takes as input a single image of the animal and produces as output a set of confidence maps (probability distributions) which describe the location of each body part within the input (**Fig. 1c**). The global maximum in each confidence map represents the network’s prediction of that body part’s position (**Fig. 1c, insets**). We employ a fully convolutional network architecture. This type of neural network eschews fully connected layers in lieu of repeated convolutions and pooling steps, which greatly improves training and prediction performance when working in the image domain ^34^.

We devised a simple 15 layer network architecture that is designed to be fast. The network consists of two blocks of 3×3×;64 convolutions, ReLU nonlinear activation, and 2-strided max pooling, which is then followed by two blocks of transposed convolutions for upsampling and additional convolutions for refinement (see **Online Methods**, **Supplementary Fig. 5a**). Pooling and downsampling allow us to keep filter sizes fixed and small, minimizing the number of computations required while allowing both local and global spatial features to be learned and combined. Recently published architectures for pose estimation follow these same general principles, but are often much larger and more complex, using skip connections, residual modules, and stacked version of the hourglass with intermediate supervision ^41^. We find that without these features, our network performs equivalently or better than those architectures (**Supplementary Fig. 5b**).

Network training consisted of a series of epochs, during which initially random weights are updated to minimize the mean-squared-error loss between ground truth and estimated confidence maps. During each epoch, 50 batches of 32 randomly sampled training images are augmented with small random rotations or reflections and evaluated for weight updates. Then, 10 batches are sampled and augmented from the held out validation set and used to compute the validation loss. This loss is used to decrease the learning rate if no significant improvements occur for multiple epochs, fine-tuning the learning process. An epoch was completed in 60 to 90 seconds on modern GPUs (see **Online Methods**).

For fast training during the labeling and initialization phase, 10% of the data are held out for validation and training is concluded after 15 epochs. After 1500 images were labeled, we proceeded to full training, for which we split the data into training (76.5%), validation (13.5%), and testing (10%) sets. We train the network for 50 epochs to increase the chance of convergence and use the held out test set to evaluate the final accuracy. All accuracy measures reported here were computed from this held out test set.

#### (iii) Pose estimation

After amortizing for initialization (loading the network onto the GPU), we find that the network is able to generate predictions at speeds suitable even for real time processing: 185±1.1 Hz (mean+-s.d.) for 192×192 images. Without any further refinement, poses generated by the network faithfully represented many features of *Drosophila* behavior that have been difficult to track automatically due to issues of occlusion, e.g., thin body parts, such as the legs, being occluded by the body or wings (**Fig. 1e**, **Supplementary Movie 1-3)**. For example, we found that the network was able to continuously and accurately track the motion of all 6 legs during extended bouts of locomotion (**Fig. 1d, Supplementary Movie 1,2**). In addition, the network can accurately track bouts of head grooming during which the forelegs are highly occluded by the head (**Fig. 1e, Supplementary Movie 3**).

### Performance of LEAP: Accuracy, speed, and training sample size

We evaluated the accuracy of LEAP after full training with 1,500 labeled images by measuring error as the Euclidean distance between estimated and ground truth coordinates of each body part on a held-out test set of 300 frames. We found that the accuracy level depends on the body part being tracked, with parts that are more often occluded, such as hind legs, resulting in slightly higher error rates (**Fig. 2a**). Overall, we found that error distances for all body parts were well below 3 pixels for the vast majority of tested images (**Fig. 2b**). This error is achieved rather quickly during training, requiring as few as 15 epochs (15-20 minutes of training time) to achieve approximately 1.97 pixel overall accuracy, and less than 50 epochs (50-75 minutes) for convergence to 1.63 pixel accuracy with the full training set (**Fig. 2c**). To measure the ground truth accuracy during the alternating labeling-training phase, we also measured the errors on the full test set as a function of the number of labeled images used for training under the fast training regime (15 epochs). We found that with as few as 10 labeled images the network is able to achieve <2.5 pixel error (2-3% of body length) in 74% of the test set, while 1,000 labeled images yields an accuracy of <2.5 pixels in 87% of the test set (**Fig. 2d**). This level of accuracy when training for few epochs with few samples contributes to the drastic reduction in time spent hand-labeling after fast training (**Supplementary Fig. 4**).

**Figure 2:**
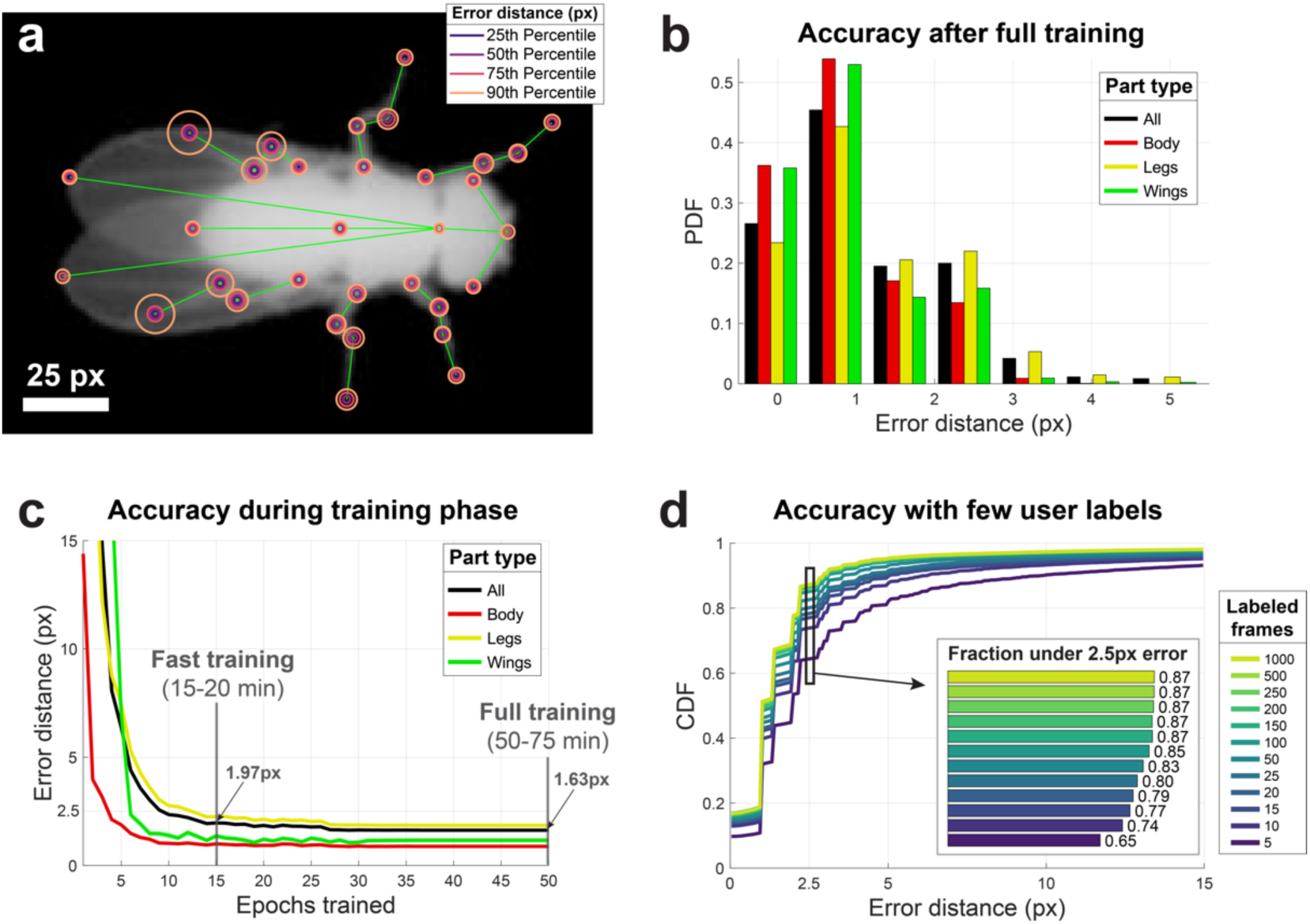
LEAP is highly accurate, and requires little training or labeled data. (a) Part-wise accuracy distribution after full training. Circles are plotted on a reference image to indicate the fraction of held out testing data (n = 300 images) for which estimated positions of the particular body part are closer to the ground truth than the radii. Most body parts have error rates below 3 pixels for over 90% of tested images. Body parts that often suffer from occlusion (e.g., hind legs) have higher rates of error. (b) Accuracy summary on held out test set after full training. Both total and grouped error rates fall well below 3 pixels (1/64th of 192×192 pixel images) in terms of Euclidean distance to ground truth as in (a). (c) Accuracy as a function of training time demonstrates fast convergence and time/accuracy trade-off during training. In the “fast training” regime, the training procedure runs for only 15 epochs, allowing the network to approximate convergence-level accuracy in a fraction of the time, optimal for training for initialization with few samples. For these tests, n = 1215 labeled frames were used for training. Lines and shaded area indicate mean and SEM for all held out test images pooled over 5 runs. After 50 epochs, convergence is achieved at the cost of additional run time. Run times depend mainly on the performance of the hardware being used, with a range provided by estimates from high end consumer or enterprise GPUs. (d) Accuracy as a function of number of training examples demonstrates the trade-off between estimation accuracy and time spent labeling. Distributions indicate estimation errors in a held out test set (n = 300 frames) while varying the number of labeled images used for training, pooled over 5 “fast training” runs. Using as few as 10 labeled images, 74% of body part estimates fell within 2.5 pixels of their ground truth locations, increasing to 87% with 1000 labeled images (inset).

### Leg tracking with LEAP recapitulates previously described gait structure

To evaluate the usefulness of our pose estimator for producing experimentally valid measurements, we used it to analyze the gait dynamics of freely moving flies. Previous work on *Drosophila* gait relied on imaging systems that use a combination of optical touch sensors and high speed video recording to follow fly legs as they walk ^24^. Although this system can accurately track fly footprints over a few seconds at a time, it cannot track the limbs when they are not in contact with the surface (during swing). Other methods to investigate gait dynamics use a semi-automated approach to label fly limbs ^26,42^. This requires a large time investment to manually correct automatically generated predictions, and therefore the semi-automated approach typically involves smaller datasets.

We began by evaluating our network on the dataset of 59 adult male fruit flies ^10^ and extracting the predicted positions of each leg tip in each of 21 million frames. For every frame in which the fly was moving forward (7.2 hours/2.6 million frames total), we encoded each leg as either in swing or stance depending on whether the leg was moving forward or backward relative to the fly’s direction of motion (**Fig. 3a**). Using this encoding, we measured the relationship between the fly’s speed and the duration of stance and swing (**Fig. 3b**). Similar to previous work, we find that swing duration is relatively constant across walking speeds, whereas stance duration decreases with walking speed ^24^. Because our methods allow us to estimate animal pose during both stance and swing (versus only during stance ^24^), we have the opportunity to investigate the dynamics of leg motion during the swing phase. We found that swing velocity increases with body speed, corroborating previous results (**Fig. 3c**). We also found that fly leg velocities follow a parabolic trajectory parametrized by body speed (**Fig. 3c**).

**Figure 3:**
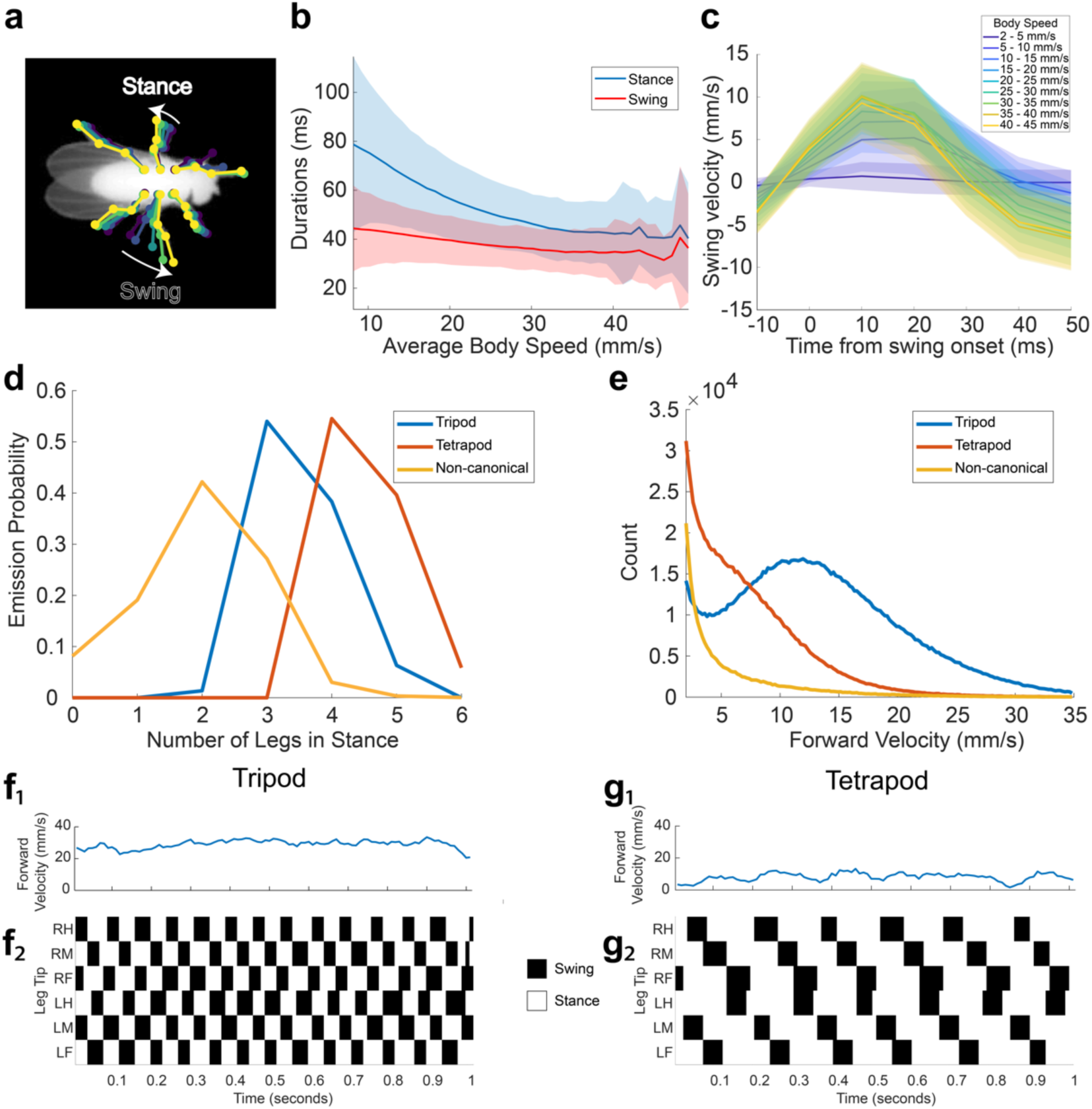
LEAP recapitulates known gait patterning in flies. (a) Schematic of swing and stance encoding. (b) Duration of swing and stance as a function of average body speed. Stance duration decreases with increasing body speed, corroborating previous findings (Mendes et al. 2013). This data comprises approximately 7.2 hours in which the fly is moving forward (2.6 million frames). Shaded regions indicate one standard deviation. (c) Swing velocity as a function of time from swing onset, and binned by body speed. Shaded regions indicate one standard deviation. (d) Emission probabilities of numbers of legs in stance for each hidden state in the HMM (see Methods). Hidden state emissions resemble tripod, tetrapod, and non-canonical gaits. (e) Distributions of velocities for each hidden state. Flies primarily exhibit tripod gait at high velocities, and tetrapod or non-canonical gaits at intermediate and slow velocities. (f,g) Examples of tripod and tetrapod gaits identified by the HMM.

Following the work of ^42^, we then trained a 3 state Hidden Markov Model (HMM) to capture the different gait modes exhibited by *Drosophila*. The emission probabilities from the model of the resulting hidden states were indicative of tripod, tetrapod, and non-canonical/wave gaits (**Fig. 3d**). As expected, we observed tripod gait at high body velocities and tetrapod or non-canonical gaits at intermediate and low velocities, in accordance with previous work ^24,42,43^ (**Fig. 3e-g**). These results demonstrate that our pose estimator is able to effectively capture the dynamics of known complex behaviors, such as locomotion.

### Body dynamics reveal structure in the fly behavioral repertoire

We next used the output of LEAP as the first step in an unsupervised analysis of the fly behavioral repertoire ^10^. We calculated the position of each body part relative to the center of the fly abdomen for each point in time and then computed a spectrogram for each of these timeseries via the Continuous Wavelet Transform (CWT). We then concatenated these spectrograms and embedded the resulting feature vectors into a two-dimensional space of actions we term a behavior space (**Online Methods, Fig. 4a**). As has been shown previously, the distribution of time points in this space is concentrated into a number of strong peaks that represent stereotyped behaviors seen across time and in multiple individuals ^10^.

**Figure 4:**
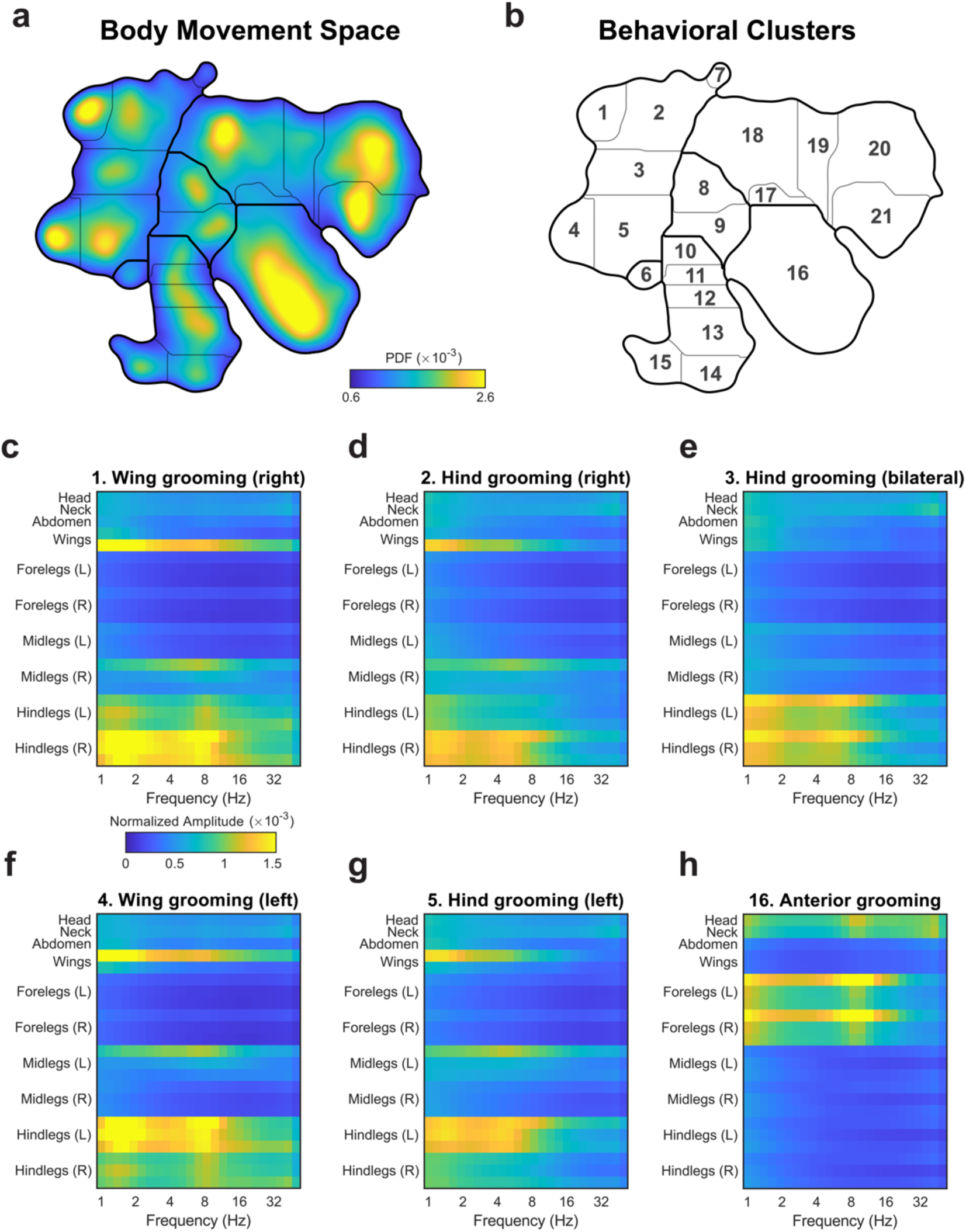
Unsupervised embedding of body position dynamics. (a) Density of freely moving fly body part trajectories, after projecting their spectrograms into to two dimensions via unsupervised nonlinear manifold embedding (Berman et al., 2014). The distribution shown is generated from 21.1 million frames. Regions in the space with higher density correspond to stereotyped movement patterns, whereas low density regions form natural divisions between distinct dynamics. A watershed algorithm is used to separate the peaks in the probability distribution (see Methods). (b) Cluster boundaries from (a) with cluster numbers indicated. (c-h) Average spectrograms from time points that fall within the dominant grooming clusters; cluster numbers are indicated in (b). Posterior grooming behaviors subdivide into symmetric clusters corresponding to the lateralization of limbs employed (c-g). Qualitative labels for each cluster based on visual inspection are provided for convenience. Colormap corresponds to normalized power for each body part.

We identify clusters in the behavior space distribution by grouping together regions of high occupancy and stereotypy (**Fig. 4b**). This distribution is sharper than what we found previously using a PCA-based compression of the images (**Supplementary Fig. 6**), with many of the least resolved behaviors now grouped together appropriately. An additional advantage to using pose estimation over PCA-based image compression is the ability to describe stereotyped behaviors by the dynamics of each body part. We calculated the average concatenated spectrogram for each cluster and found that specific behaviors are recapitulated in the motion power spectrum for each body part (**Fig. 4c-h**).

This method can be used to accurately describe grooming, a class of behaviors that is highly represented in our dataset. Posterior grooming behaviors exhibited a distinctly symmetric topology (**Fig. 4b-g**), revealing both bilateral (**Fig. 4e**) as well as unilateral grooming of the wings (**Fig. 4c,f**) and the rear of the abdomen (**Fig. 4d,g**). These behaviors involve unilateral, broadband (1-8 Hz) motion of the hind legs on one side of the body and a slower (∼1.5 Hz) folding of the wing on the same side of the body. In contrast, anterior grooming is characterized by broadband motions of both front legs with a peak at ∼9 Hz, representing the legs rubbing against each other (**Fig. 4h**).

We also discovered a number of unique clusters related to locomotion (**Fig 5a,b**). The slowest state (cluster 10) involves a number of frequencies with a broad peak centered at 5.1 Hz (**Fig. 5 c-e**). This can be seen both in the concatenated spectrograms (**Fig. 5c**) and the power spectrum averaged over all leg positions (**Fig. 5d**). The fly center-of-mass velocity distribution for this behavior is shown in **Figure 5e**. As the fly speeds up (clusters 10-15, **Fig. 5e**), the peak frequency for the legs increases monotonically to 11.5 Hz (cluster 15). We next asked if the tripod and tetrapod gaits we found in our previous analysis were represented by distinct regions in the behavior space. We found that tripod gait was used predominantly in the three fastest locomotion behaviors whereas the tetrapod (and to a lesser extent the non-canonical) gait was used for the three slower locomotion behaviors (**Supplementary Fig. 5f**).

**Figure 5:**
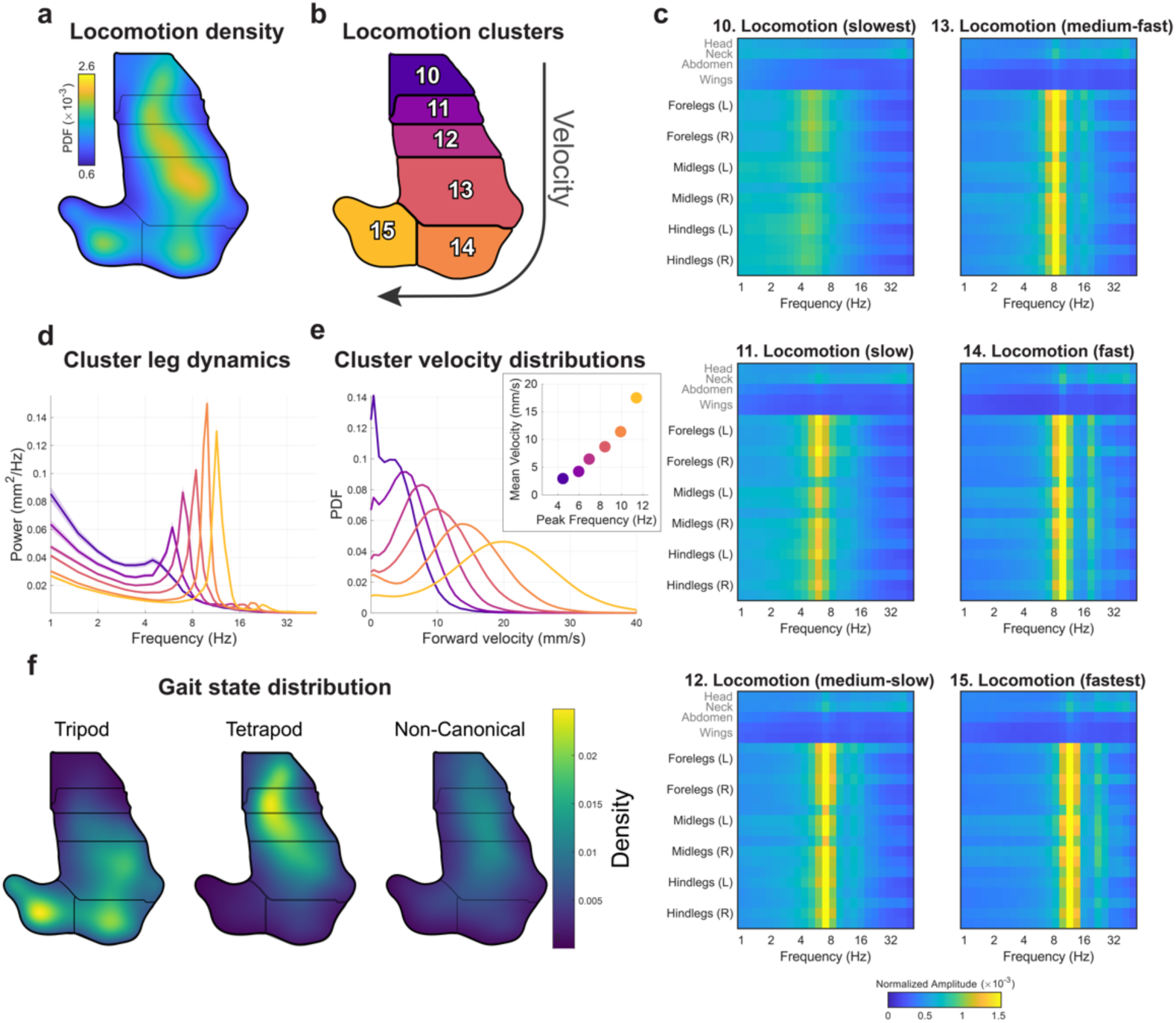
Locomotor clusters in behavior space separate distinct gait modes. (a, b) Density and cluster labels of locomotion clusters (from the same behavioral space shown in Fig. 4a). (c) Average spectrograms (similar to Fig. 4c-h) quantify the dynamics in each cluster. The frequency spectrum of leg movements in each cluster is sharp and shifts from 5.1 to 11.5 Hz from slowest to fastest locomotion speeds. (d) Average power spectra calculated from the leg joint positions for each cluster in (c). Colors correspond to the cluster numbers in (b). Each spectrum has a single dominant peak between 5.1 and 11.5 Hz, with harmonics from 12-25 Hz seen in the fastest subtypes. (e) The distribution of forward locomotion velocity exhibits a peak that shifts to the right as a function of cluster number. Colors correspond to cluster numbers in (b). (inset) Forward locomotion velocity increases with peak leg frequency. (f) Gait modes identified by HMM from swing/stance state correspond to distinct clusters.

### LEAP generalizes to images with complex backgrounds or of other animals

To test the robustness and generalizability of our approach under more varied imaging conditions, we evaluated the performance of LEAP on a dataset in which pair of flies were imaged against a non-uniform and low contrast background of porous mesh (∼4.2 million frames, ∼11.7 hours) (**Fig. 6a_1_**). Using the same workflow as in the first dataset, we found that the pose estimator was able to reliably recover body part positions with high accuracy despite poorer illumination and a complex background that was at times indistinguishable from the fly (**Fig. 6a_2,3_, Supplementary Movie 4**). We then applied a previously described method for segmentation and tracking ^13^ to these images to evaluate the performance when masking out the background (**Fig. 6b_1_**). Even with substantial errors in the masking (e.g., leg or wing segmentation artifacts), we find that the accuracy remains high and is improved slightly by excluding the background pixels from the images when compared to the raw images (**Fig. 6b_2,3_, Supplementary Movie 4**). Finally, we tested the applicability of our framework to animals with different morphology by tracking videos of freely behaving mice (*Mus musculus*) imaged from below in an open arena (**Fig. 6c_1_**). We observed comparable accuracy in these mice despite considerable occlusion during behaviors such as rearing (**Fig. 6c_2,3_, Supplementary Movie 5**).

**Figure 6:**
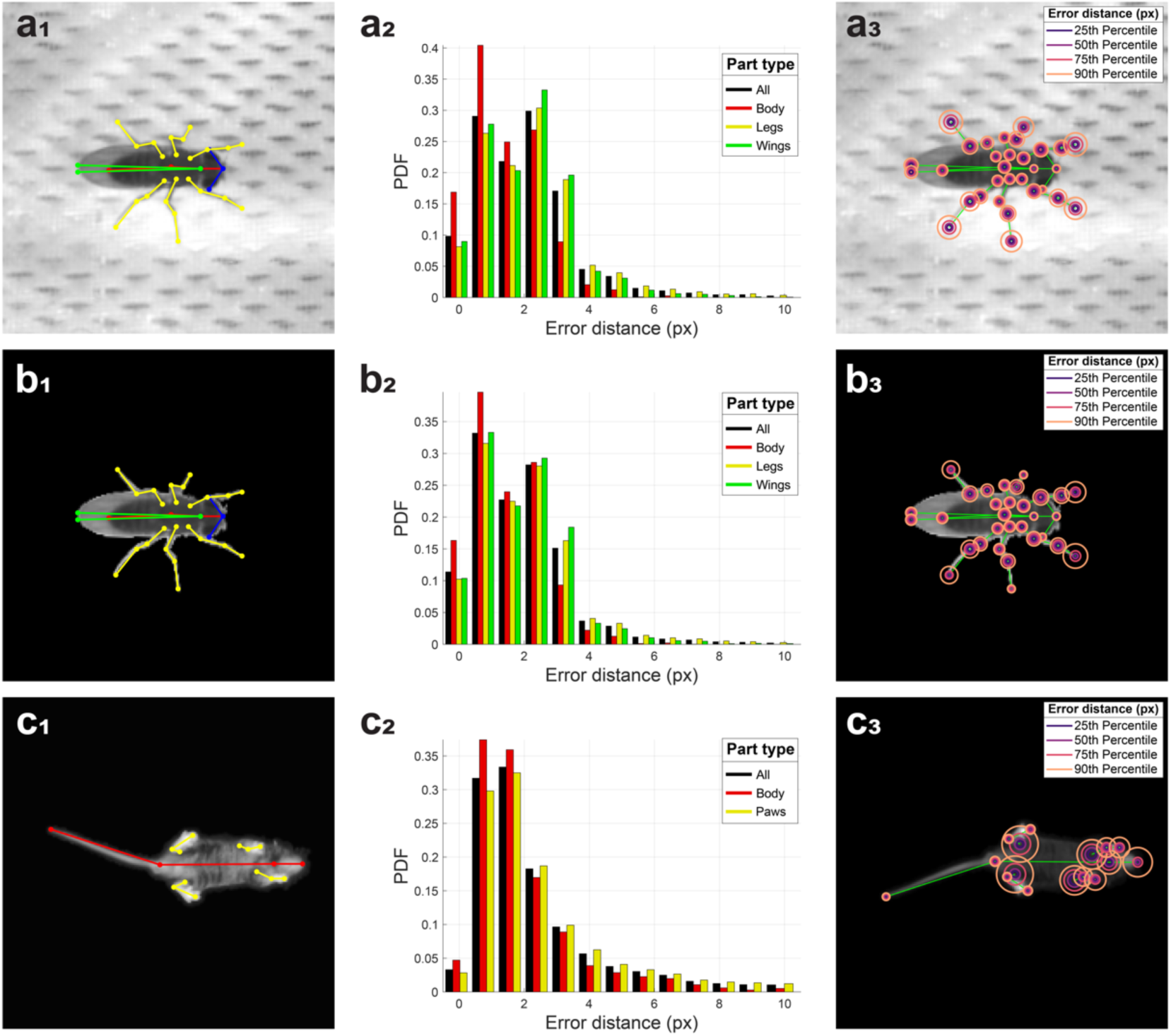
LEAP generalizes to images with complex backgrounds or of other animals. (a) LEAP estimates on a separate dataset of 42 freely moving male flies, each imaged against a heterogeneous background of mesh and microphones, with side illumination (∼4.2 million frames, ∼11.7 hours). 32 body parts (see Supp Fig. 3) were tracked (a_1_), and 1,530 labeled frames were used for training. Error rates for position estimates were calculated on a held out test set of 400 frames (a_2_) and were comparable to those achieved for images with higher signal to noise (compare with Fig. 2b). Part-wise error distances (a_3_) illustrate that accuracy is lower in distal body parts, likely due to ambiguity with the background mesh holes. (b) LEAP estimates on masked images from the dataset described in (a). Background was subtracted using standard image processing algorithms (see Methods) to reduce the effect of background artifacts. Similar accuracy measures are observed (compare b_2_ with a_2_). Error distances are higher for distal body parts that are often masked out due to the difficulty in resolving those pixels from the background (b_3_). (c) LEAP estimates on a dataset of freely moving mice imaged from below (∼3 million frames, ∼4.8 hours). Three points are tracked per leg, in addition to the tip of the snout, neck, and base and tip of the tail (c_1_) - 1000 labeled frames were used for training. Accuracy rates on a held out test set (of 242 frames) are higher but still comparable to fly datasets (c_2_). Most errors come from the leg base point, which is often occluded (c_3_).

## Discussion

Here we present a pipeline (termed LEAP) that uses a deep neural network to track the body parts of a behaving animal in all frames of a movie via labeling of a small number of images from across the dataset. We show that this method is fast (requiring one hour to train and producing body part position estimates at a maximum rate of 185 Hz), accurate (training on 10 frames results in 74% of estimates within 2.5 pixel error while training on 100 frames results in 85% of the frames within 2.5 pixel error), and generalizes across animal species (including flies and mice) and different regimes of signal to noise ratio. Importantly, we do not construct a single network to perform pose estimation on all datasets, but rather we present a single architecture that can be trained to perform pose estimation on any dataset if given a small number of training samples. All that is required of future users is that the training sets be compiled in a specific manner that can be facilitated with our user interface (for which we provide code and utilities).

Discovering the proximate mechanisms underlying behavior relies on an analysis of behavioral dynamics matched to the timescales of neural and muscular activity. Tracking only the centroid of an animal and its change in position or heading over time is likely an insufficient level of description for determining how the nervous system controls most behaviors. Previous studies have addressed the issue of pose estimation either through centroid tracking ^2^, pixel-wise correlations ^10,11^, or specialized apparatus for tracking body parts ^17,20,24,42,44^. For the latter, applying markers to an animal can limit natural behavior and systems that track particular body parts are not in general scalable to all body parts or animals with a very different body plan. We demonstrate the value of LEAP by showing how it can be applied to the study of locomotor gait dynamics (**Fig. 3, 5**) and unsupervised behavioral mapping (**Fig. 4, 5**) in *Drosophila*. Previous studies of gait dynamics have been limited to short stretches of locomotor bouts that were captured using a specialized imaging system ^24^ or to the number of behavioral frames that could be hand-labeled ^42^. We show that LEAP not only recapitulates previous findings on locomotor gait, but that it also discovers new aspects of the behavior (for example, that the dynamics of the leg during swing have a nonlinear relationship with swing velocity). We also demonstrate the clear interpretability afforded when using LEAP in combination with unsupervised behavior classification (**Fig. 4, 5**). This provides a solution to a major shortcoming in existing approaches, namely that identified behaviors had to be interpreted simply by watching videos ^10,11^. Using LEAP as the first step in such unsupervised algorithms, each discovered behavior can now be interpreted by analyzing the dynamics of each body part.

There are a number of applications for this pipeline beyond those demonstrated here. Because the network learns body positions from a small amount of human labeled frames, the network can be easily trained to track a wide variety of animal species and classes of behavior. Further, LEAP can be extended to tracking of body parts in 3D by either using multiple cameras or depth-sensitive devices. This will likely be useful for tracking body parts of head-fixed animals moving on an air supported treadmill ^45,46^. These experiments are particularly suited for our approach, as the movies from head-fixed animals are inherently recorded in egocentric coordinates. Additionally, we note that the fast prediction performance of our method makes it compatible with closed-loop experimentation, where joint positions may be computed in realtime to control experimental parameters such as stimuli presented to the animal or optogenetic modulation. Lastly, through the addition of a segmentation step for analyzing movies of multiple animals ^2,13,39^, LEAP can estimate poses for multiple interacting individuals.

The primary practical limitation of this framework is the egocentric alignment step that may be sensitive to imaging conditions and the context of the experiment. We note, however, that many standard techniques exist to find the centroid and orientation of animals in images, including deep learning-based approaches ^40^. Other concerns may pertain to generalizability, in particular due to how we train each network from scratch rather than performing transfer learning to reuse a set of more general, shallow layer feature detectors ^36^. While transfer learning could easily be incorporated into LEAP (as well as any other network architecture designed for pose estimation), we found it to be unnecessary given the inherently low variability of imaging conditions in the lab and the empirically determined low training data requirements.

In summary, we present a method for tracking body part positions of freely moving animals with little manual effort and without the use of physical markers. We show LEAP’s robustness, state-of-the-art performance, validity, and utility for quantitative behavioral analysis. We anticipate that this tool will reduce the technical barriers to addressing a broad range of previously intractable questions in ethology and neuroscience through quantitative analysis of the dynamic changes in the full pose of an animal over time.

## Contributions

Designed study: TP, DA, SW, JS, and MM

Conducted experiments: TP, DA, LW, and MK

Developed GUI and analyzed data: TP and DA

Wrote manuscript: TP, DA, JS, and MM

## Acknowledgments

The authors acknowledge Jonathan Pillow for helpful discussions; Gordon Berman and Daniel Choi for providing fly behavior data; Byung Cheol Cho for contributions to the mouse experimental setup, acquisition and preprocessing pipeline; Peter Chen for a previous version of a neural network for pose estimation that was useful in designing our method; Heejae Jang, Malavika Murugan, and Ilana Witten for feedback on the GUI and other helpful discussions; Georgia Guan for assistance maintaining flies; and the Murthy, Shaevitz and Wang labs for general feedback.

This work was supported by the NIH: R01 NS104899-01 BRAIN Initiative Award (to MM and JS), R01 MH115750 (to SW and JS), NIH R01 NS045193; NSF BRAIN Initiative EAGER Award (to MM and JS); the Nancy Lurie Marks Family Foundation (to SW); an HHMI Faculty Scholar Award (to MM); and an NSF GRFP DGE-1148900 (to TP).

## Methods

### Code

The code for running LEAP, as well as all accompanying GUIs, trained networks, labeled data and analysis code for figure reproduction, can be found in the following repository: https://github.com/talmo/leap

### Datasets

Details on the dataset of 59 adult male Drosophila can be found in ^1,2^. Animals were allowed to move freely in a backlit 100mm diameter circular arena covered by a 2mm tall clear PETG dome. Videos were captured from the top with a Point Grey Gazelle camera at a resolution of ∼35 pixels/mm at 100 FPS for 1 hour for each fly, totaling ∼21 million frames for the dataset. To calculate the spatial resolution of LEAP we assumed a mean male fly length of 2.82mm ^3^.

The second fly dataset reported here (**Fig. 5**) consists of 42 videos of freely moving pairs of virgin male and female fruit flies (NM91 strain), 3-5 days post-eclosion. Only males from these videos were analyzed in this study. Flies moved freely within a 30mm diameter circular arena with a 2mm tall clear PETG dome against a white mesh floor covering an array of microphones, resulting in an inhomogeneous image background. Videos were captured from above using a Point Grey Flea3 camera at a resolution of ∼25 pixels/mm at 100 FPS, totaling ∼4.2 million frames.

The mouse dataset for **Figure 5** consisted of 29 videos of C57/BL6 strain mice (*Mus musculus*), 15 weeks (108 days) old. Animals moved freely in a 45.7×45.7 cm open field arena with a clear acrylic floor for 10 minutes each. Videos were captured from below with IR illumination using a Point Grey Blackfly S camera at a resolution of 1.95 pixels/mm at 170 FPS, totaling ∼3 million frames. Experimental procedures were approved by the Princeton University Institutional Animal Care and Use Committee and conducted in accordance to the National Institutes of Health guidelines for the humane care and use of laboratory animals. Mice used in this study were ordered through Jackson Laboratory (The Jackson Laboratory, Bar Harbor, ME) and had at least one week of acclimation to the Princeton Neuroscience Institute vivarium before experimental procedures were performed. Mice were kept in group cages with food and water ad libitum under a reversed 12:12 hour dark-light cycle (light: 19:30-7:30).

### Preprocessing and alignment to generate egocentric images for labeling and training in LEAP

For the main fly dataset (59 males), we used the alignment algorithm from ^1^. The raw videos consisted of unoriented bounding boxes around the flies from a closed-loop camera tracking system. Individual frames were then aligned to a template image of an oriented fly by matching the peak of the Radon transformed fly image to recover the orientation and then computing the cross correlation to center the fly. The centroid and orientation parameters were used to crop a 200×200 pixel oriented bounding box in each frame. Code for alignment is available in the repository accompanying the original paper: https://github.com/gordonberman/MotionMapper

For the second fly dataset (42 males), we adapted a previously published method for tracking and segmentation of videos of courting fruit flies ^4^. We first modeled the mesh background of the images by fitting a normal distribution to each pixel in the frame across time with a constant variance to account for camera shot noise. The posterior was evaluated at each pixel of each frame and then thresholded to segment the foreground pixels. Due to the inhomogeneity of the arena floor mesh, significant segmentation artifacts were introduced, particularly when translucent or very thin body parts (i.e., wings and legs) could not be disambiguated from the dark background mesh holes. The subsequent steps of histogram thresholding, morphological filtering and ellipse fitting were performed as described previously in ^4^. We developed a simple GUI for proofreading the automated ellipse tracking before extracting 200 ×; 200 pixel oriented bounding boxes. We extracted bounding boxes for both animals in each frame and saved both the raw pixels containing the background mesh as well as the foreground-only images which contain segmentation artifacts. This pipeline was implemented in MATLAB and the code is available in the code repository accompanying this paper.

For the mouse videos, a separate preprocessing pipeline was developed. Raw videos were processed in three stages: (1) animal tracking, (2) segmentation from background, and (3) alignment to the body centroid and tail-body interface. In stage (1), the mouse’s torso centroid was tracked by subtracting a background image (median calculated at each pixel value across that video), retrieving pixels with a brightness above a chosen threshold from background (mice were brighter than background), and using morphological opening to eliminate noise and the mouse’s appendages. The largest contiguous region reliably captured the mouse’s torso (referred to below as the torso mask) and was used to fit an ellipse whose center was used to approximate the center of the animal. In stage (2), a similar procedure as in stage (1) was employed to retrieve a full body mask. In this stage, a more permissive threshold and smaller morphological opening radius were used than in stage (1) to capture the mouse’s body edges, limbs, and tail while still eliminating noise. The pixels outside of this body mask were set to 0. In stage (3) each segmented video frame was translated and rotated such that frame’s center coincided with the center of the animal and the x-axis lay on the line connecting the center and tail-body attachment point. The tail-body attachment point was defined as the center of a region overlapping between the torso mask and a dilated tail mask. The tail mask was defined as the largest region remaining after subtracting the torso mask from the full body mask and performing a morphological opening. After applying these masks to segment the raw images, bounding boxes were extracted by using the ellipse center and orientation.

Oriented bounding boxes were cropped to 192 ×; 192 pixels for all datasets to ensure consistency in output image size after repeated pooling and upsampling steps in the neural network. These data were stored in self-describing HDF5 files.

### Sampling diverse images for labeling and training in LEAP

To ensure diversity in image and pose space when operating at low sample sizes, we employ a multistage cluster sampling technique. First, *n*_*0*_ images were sampled uniformly from each dataset by using a fixed stride over time to minimize correlations being temporally adjacent samples. We then used principal component analysis (PCA) to reduce their dimensionality, and the images were then projected down to the first *D* principal components. After dimensionality reduction, the images were grouped via k-means clustering *k* into subgroups from which *n* images were randomly sampled from each group. To minimize the time necessary for the network to generalize to images from all groups, we sorted the dataset such that consecutive samples cycled through the groups. This way, uniform sampling was maintained even at the early phases of user labeling, ensuring that even a network trained on only the first few images will be optimized to estimate body part positions for a diversity of poses. We used *n*_0_= 500, yielding 29,500 initial samples; *D* = 50, which is sufficient to explain 80% of the variance in the data (**Supplementary Fig. 2**); *k*= 10and *n* = 150 to produce a final dataset of 1,500 frames for labeling and training.

### LEAP neural network design and implementation

We based our network architecture on previous designs of neural networks for human pose estimation ^5–7^. We adopt a fully convolutional architecture that learns a mapping from raw images to a set of confidence maps. These maps are images that can be interpreted as the 2-d probability distribution (i.e., heatmap) centered at the spatial coordinates of each body part within the image. We train the network to output one confidence map per body part stacked along the channel axis.

Our network consists of 15 layers of repeated convolutions and pooling (**Supplementary Fig. 4**). The convolution block consists of 3x convolution layers (64 filters, 3×3 kernel size, 1×1 stride, ReLU activation). The full network consists of 1x convolution block, 1x max pooling across channels (2×2 pooling size, 2×2 stride), 1x convolution block (128 filters), 1x max pooling (2×2 pooling size, 2×2 stride), 1x convolution block (256 filters), 1x transposed convolution (128 filters, 3×3 kernel size, 2×2 stride, ReLU activation, Glorot normal initialization), 2x convolution (128 filters, 3×3 kernel size, 1×1 stride, ReLU activation), and 1x transposed convolution (128 filters, 3×3 kernel size, 2×2 stride, linear activation, Glorot normal initialization).

We base our decisions of these hyperparameters on the idea that repeated convolutions and strided max pooling enable the network to learn feature detectors across spatial scales. This allows the network to learn how to estimate confidence maps using global image structure which provides contextual information that can be used to improve estimates even for occluded parts ^5,7^. Despite the loss of resolution from pooling, the upsampling learned through transposed convolutions is sufficient to recover the spatial precision in the confidence maps. We do not employ skip connections, residual modules, stacked networks, regression networks, or affinity fields in our architecture as used in other approaches of human pose estimation ^5,6,8,9^.

For comparison, we also implemented the stacked hourglass network ^7^. We tested both the single hourglass version and 2x stacked hourglass with intermediate supervision. The hourglass network consisted of 4x residual bottleneck modules (64 output filters) with max pooling (2×2 pool, 2×2 stride), followed by their symmetric upsampling blocks and respective skip connections. The stacked version adds intermediate supervision in the form of a loss term on the output of the first network in addition to the final output.

We implemented all versions of neural networks in Python via Keras and TensorFlow, popular deep learning packages that allow transparent GPU acceleration and easy portability across operating systems and platforms. All Python code was written for Python 3.6.4. Required libraries were installed via the pip package manager: numpy (1.14.1), h5py (2.7.1), tensorflow-gpu (1.6.0), keras (2.1.4). We tested our code on machines running either Windows 10 (v1709) and a RedHat-based Linux distribution (Springdale 7.4) with no additional steps required to port the software other than installing the required libraries.

Code for all network implementations is available in the main repository accompanying this paper.

### LEAP training procedure

Prior to training, we generated an augmented dataset from the user-provided labels and corresponding images. We first doubled the number of images by mirroring the images along the body symmetric axis and adjusting the body part coordinates accordingly, including swapping left/right body part labels (e.g., legs). Then, we generated confidence maps for each body part in each image by rendering the 2-d Gaussian probability distribution centered at the ground truth body part coordinates, *μ* = (x,y), and fixed covariance, ∑= *diag*(*σ*) with a constant *σ* = 5*px*. These were pre-generated and cached to disk to minimize the necessary processing time during training.

Once confidence maps were computed for each image, we split the dataset into training, validation and test sets. The training set was used for backpropagation of the loss for updating network weights, the validation set was used to estimate performance and adjust the learning rate over epochs, and the test set was held out for analysis. For the fast training, the dataset was split into only training (90%) and validation (10%) sets to make the best use of data when training with very few labels. For full training, the dataset was split into training (76.5%), validation (13.5%) and testing (10%) sets. All analyses reported here share the same held out test set to ensure it is never trained against for any replicate.

All training was done using the Adam optimizer with default parameters as described in the original paper ^10^. We started with a learning rate of 1e-3 but use a scheduler to reduce it by a factor of 0.1 when the validation loss fails to improve by a minimum threshold of 1e-5 for 3 epochs. The loss function optimized against is simply the mean squared error between estimated and ground truth confidence maps.

During training, we considered an epoch to be a set of 50 batches of 32 images, which were sampled randomly with replacement from the training set and augmented by applying a random rotation to the input image and the corresponding ground truth confidence maps. At the end of 50 batches of training, 10 batches were sampled from the separate validation set, augmented and evaluated and the loss was used for learning rate scheduling. An epoch evaluated in 60 to 90 seconds including all augmentation, forward and reverse passes, and the validation forward pass when running on a modern GPU (NVIDIA GeForce GTX 1080 Ti or P100).

We ran this entire procedure for 15 epochs during the fast training stage, and for 50 epochs during the full training stage. For analyses, a minimum of 5 replicates were fully trained on each dataset to estimate the stability of optimization convergence.

### Pose estimation from confidence maps

Predictions of body part positions were computed directly on the GPU. We implement a channel-wise global maximum operation to convert the confidence maps into image coordinates as a TensorFlow function, further improving runtime prediction performance by avoiding the costly transfer of large confidence map arrays. All prediction functions including normalization and saving were implemented as a self-contained Python script with a command-line interface for ease of batch processing.

### Computing hardware

All performance tests were conducted on a high end consumer-grade workstation equipped with a Intel Core i7-5960X CPU, 128 GB DDR4 RAM, NVMe SSD drives, and a single NVIDIA GeForce 1080 GTX Ti (12 GB) GPU. We also use Princeton University’s High Performance Computing cluster with nodes equipped with NVIDIA P100 GPUs for batch processing. These higher end cards afford a speed-up of ∼1.5x in the training phase.

### Accuracy analysis

For all analyses of accuracy (**Figs. 2, 6; Supplementary Figs. 4, 5**), we trained at least 5 replicates of the network with the same training/validation/testing datasets. All analyses were performed in MATLAB R2018a (MathWorks). We used the gramm toolbox for figure plotting ^11^.

### Gait analysis

We translated the body position coordinates to egocentric coordinates by subtracting the predicted location of the intersection between the thorax and abdomen from all other body position predictions for each frame. We then calculated the instantaneous velocity along the rostrocaudal axis of each leg tip within these truly egocentric reference coordinates. The speed of each body part was smoothed using a Gaussian filter with a five frame moving window. For each leg tip, instances in which the smoothed velocity was greater than zero were defined as swing while those less than zero were defined as stance. Information from this egocentric axis was combined with allocentric tracking data to incorporate speed and orientation information. The centroids and orientations of the flies were smoothed using a moving mean filter with a five frame window to find the instantaneous speed and forward velocity. To remove idle bouts and instances of backward walking, all gait analyses were limited to times when the fly was moving in the forward direction at a velocity greater than 2 mm/s (approximately one body length/s) unless otherwise noted. The analyses relating stance and swing duration to body velocity were limited to forward velocities greater than 7.2 mm/s, to remain in line with previous work ^12^.

To measure gait modes, we trained an HMM to model gait as described previously ^13^. The training data consisted of a vector denoting the number of legs in stance for bouts in which the fly was moving forward at a velocity greater than 2 mm/s lasting longer than 0.5 seconds. Training data were sampled such that up to 3,000 frames were taken from each video, resulting in a total of 159,270 frames. We trained a three-state HMM using the Baum-Welch algorithm and randomly initialized transition and emission probabilities ^14^. We designated each hidden state as tripod, tetrapod, and non-canonical in accordance with the estimated emission probabilities. We then used the Viterbi algorithm along with our estimated transition and emission matrices to predict the most likely sequence of hidden states from which the observed stance vectors for the entire dataset would emerge ^15^.

### Unsupervised embedding of body part dynamics

In order to create a map of motor behaviors described by body part movements, we used a previously described method for discovering stereotypy in postural dynamics ^1^. First, body part positions were predicted for each frame in our dataset to yield a set of 32 timeseries of egocentric trajectories in image coordinates for each video. These timeseries were recentered by subtracting the thorax coordinate at each timepoint and rescaled to comparable ranges by z-scoring each timeseries. The timeseries were then expanded into spectrograms by applying the Continuous Wavelet Transform (CWT) parametrized by the Morlet wavelet as the mother wavelet and 25 scales chosen to match dyadically spaced center frequencies spanning 1 to 50 Hz. This time-frequency representation augments the instantaneous representation of pose at each timepoint to one that captures oscillations across many timescales. The instantaneous spectral amplitudes of each body part were then concatenated into a single vector of length 2(*J* - 1)*F* where *J* is the number of body parts before subtracting the body part used as reference (i.e., the thorax) and doubled to account for both *x* and *y* coordinates, and *F* is the number of frequencies being measured via CWT. In our data, this resulted in a 1,550-dimensional representation at each timepoint.

Finally, we performed nonlinear dimensionality reduction on these high dimensional vectors by using a nonlinear manifold embedding algorithm ^16^. We first selected representative timepoints via importance sampling, wherein a random sampling of timepoints in each video is embedded into a 2D manifold via t-distributed stochastic neighbor embedding (t-SNE) and clustered via the watershed transform. This allowed us to choose a set of timepoints from each video that were representative of their local clusters, i.e., spanning the space of postural dynamics. A final behavior space distribution was then computed by embedding the selected representative timepoints using t-SNE to produce the full manifold of postural dynamics in two dimensions.

After projecting all remaining timepoints in the dataset into this manifold, we computed their 2-d distribution and smoothed with a Gaussian kernel with σ = 0.65 to approximate the probability density function of this space. We clipped the range of this density map to the range [0.5×; 10^−3^, 2.75×;^−3^] to exclude low density regions and merge very high density regions. We then clustered similar points by segmenting the space into regions of similar body part dynamics by applying the watershed transform to the density. Although both the manifold coordinates representation of each timepoint are not immediately meaningful, we were able to derive an intuitive interpretation of each cluster by referring to the high dimensional representation of their constituent timepoints. To do this, we sampled timepoints from each cluster and averaged their corresponding high dimensional feature vector, which we can then visualize by reshaping it into a body part-frequency matrix (**Fig. 4**).

## Supplemental Figures and Movie Legends

**Supplementary Figure 1:**
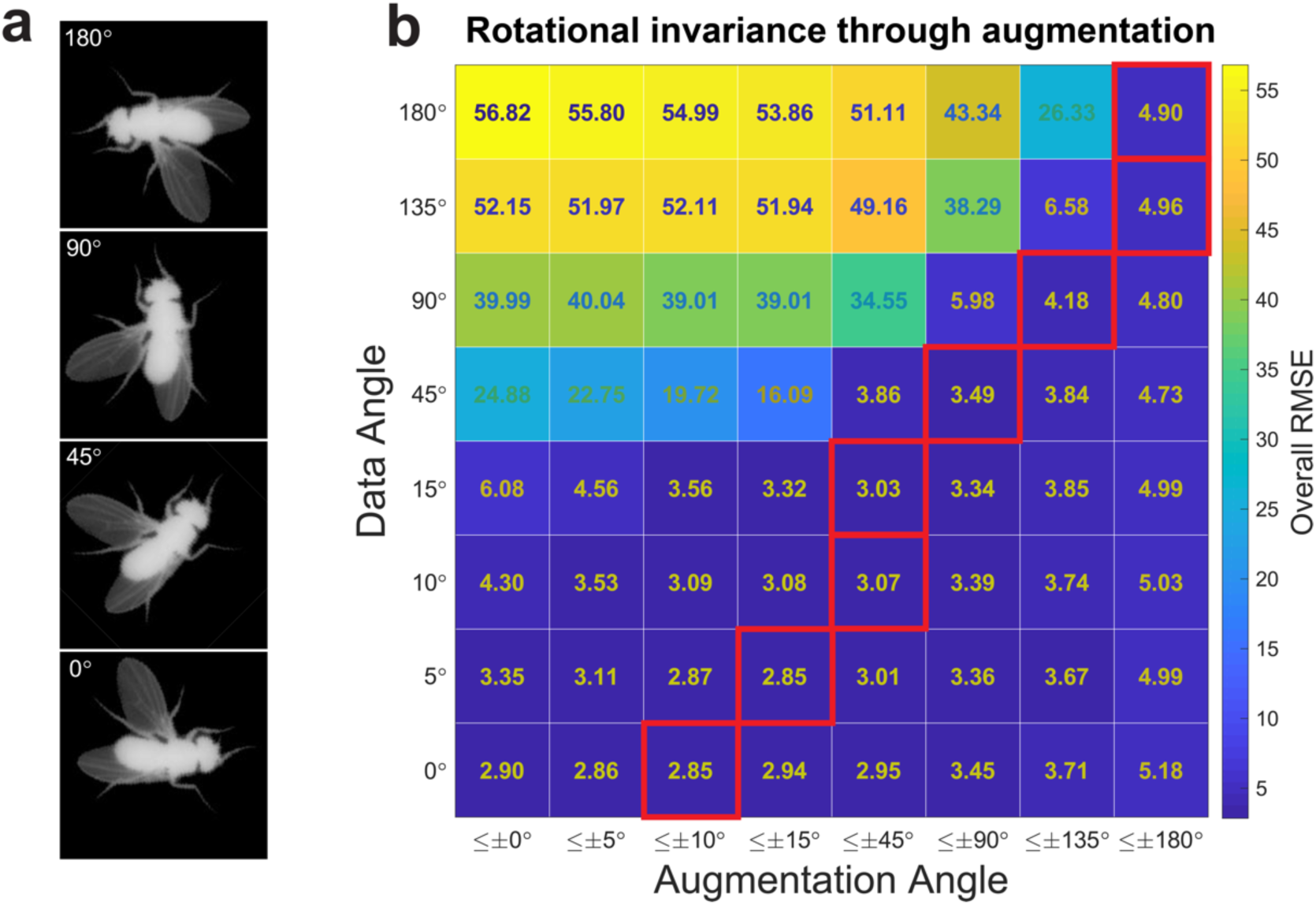
Rotational invariance is learned at the cost of prediction accuracy. (a) Rotations are applied about the center of the image. During training, confidence maps are rotated accordingly. (b) The accuracy measured as the RMSE of position estimates when evaluated on data rotated at a fixed angle (rows) with networks trained on data augmented by rotations between a range of angles (columns). Red boxes denote the best accuracy for each data angle, denoting that optimal performance is achieved when the network is trained on augmented images with the minimally inclusive range of angles. Top accuracy decreases relative to the degree of rotational invariance the network must learn.

**Supplementary Figure 2:**
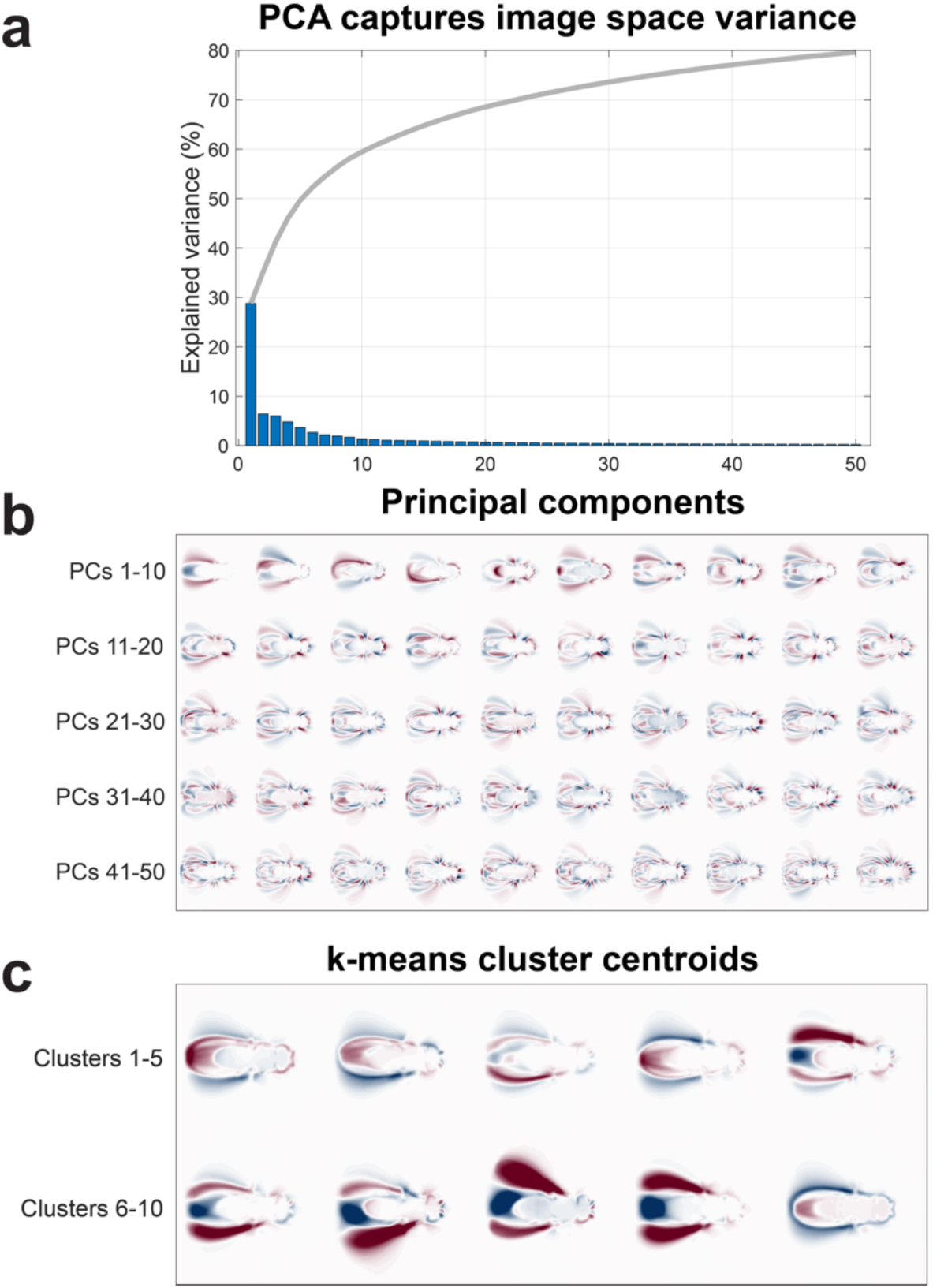
Cluster sampling to promote pose diversity in labeling dataset. (a) Principal component analysis (PCA) of unlabeled images captures the majority of the variance in the data within 50 components. The cumulative variance explained (line) suggests that using PCA for dimensionality reduction does not sacrifice substantial loss of information within the images. (b) Top PCA eigenmodes visualized as coefficient images. Red and blue shading denote positive and negative coefficients at each pixel. Areas of similar colors indicate correlated pixel intensities within a given mode. After mean subtraction, each image in the initially sampled dataset is projected onto all 50 eigenmodes. (c) Cluster centroids identified by k-means after PCA. Red and blue shading denote pixels with higher or lower intensity than the overall mean. Cluster centroids illustrate the diversity of poses that are detected in image space by this sampling method. Samples are then drawn evenly from each cluster to select representative images for labeling with the GUI.

**Supplementary Figure 3:**
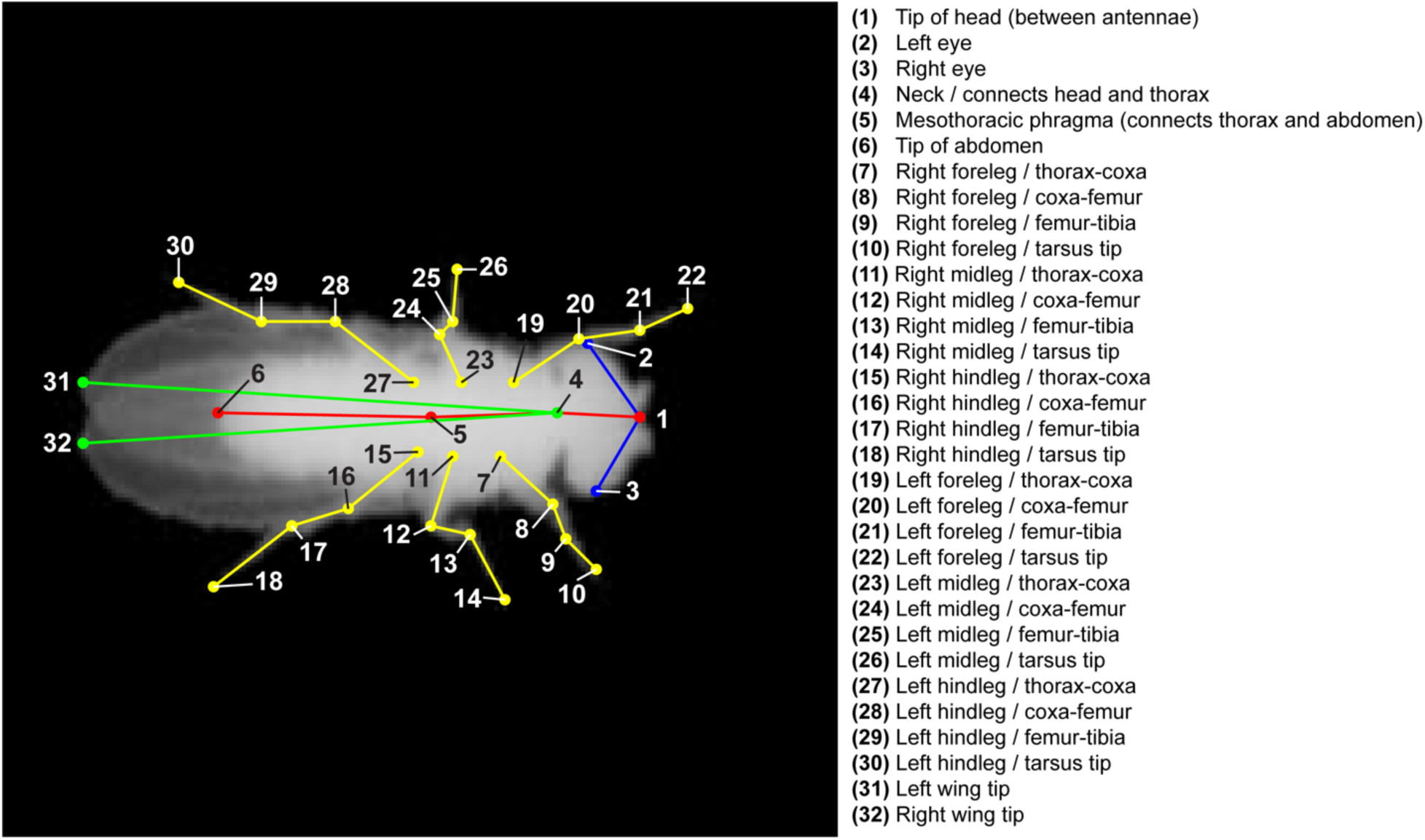
User-defined skeleton. We selected 32 points to cover the body parts of the fly; these parts were chosen to approximately match the set of visible joints and interest points in the anatomy of the animal.

**Supplementary Figure 4:**
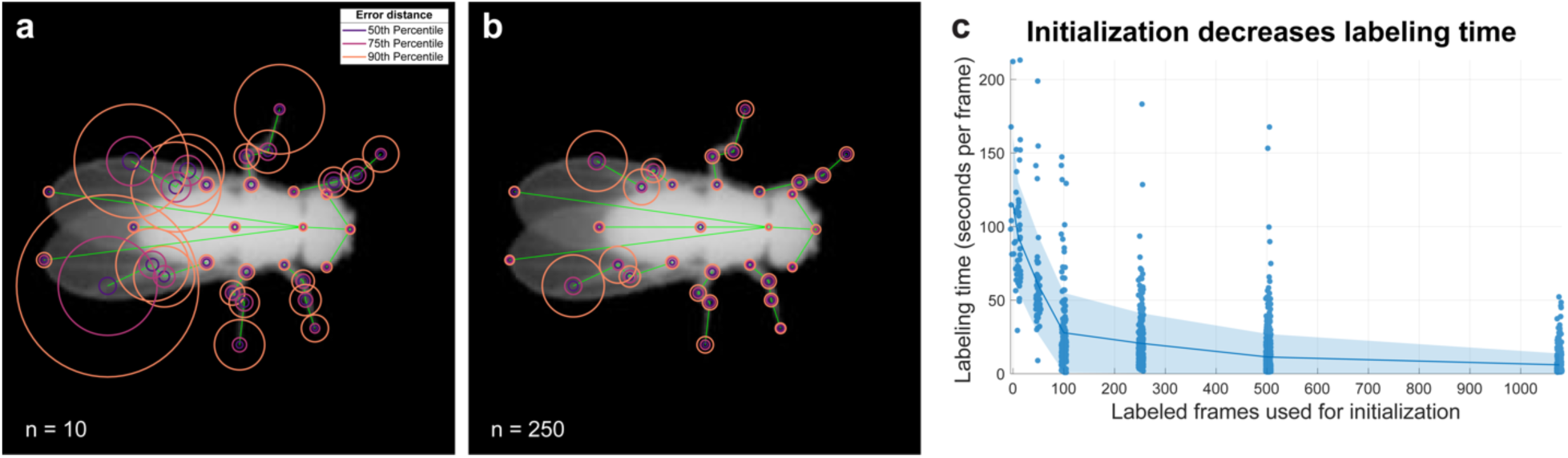
Estimation accuracy improves with few samples. (a-b) Error distance distributions per body part when estimated with networks trained for 15 epochs on 10 (a) or 250 (b) labeled frames. The majority of estimates fall within few pixels of the ground truth, reducing the labeling procedure to simply correcting estimates. (c) Time spent labeling each frame decreases with the quality of initialization. Line and shaded region correspond to mean and standard deviation. Starting frames require 115.4+-45.0 (mean+-s.d.) seconds to label, decreasing to 6.1±7.7 seconds after initializing with a network trained on 1000 labeled frames.

**Supplementary Figure 5:**
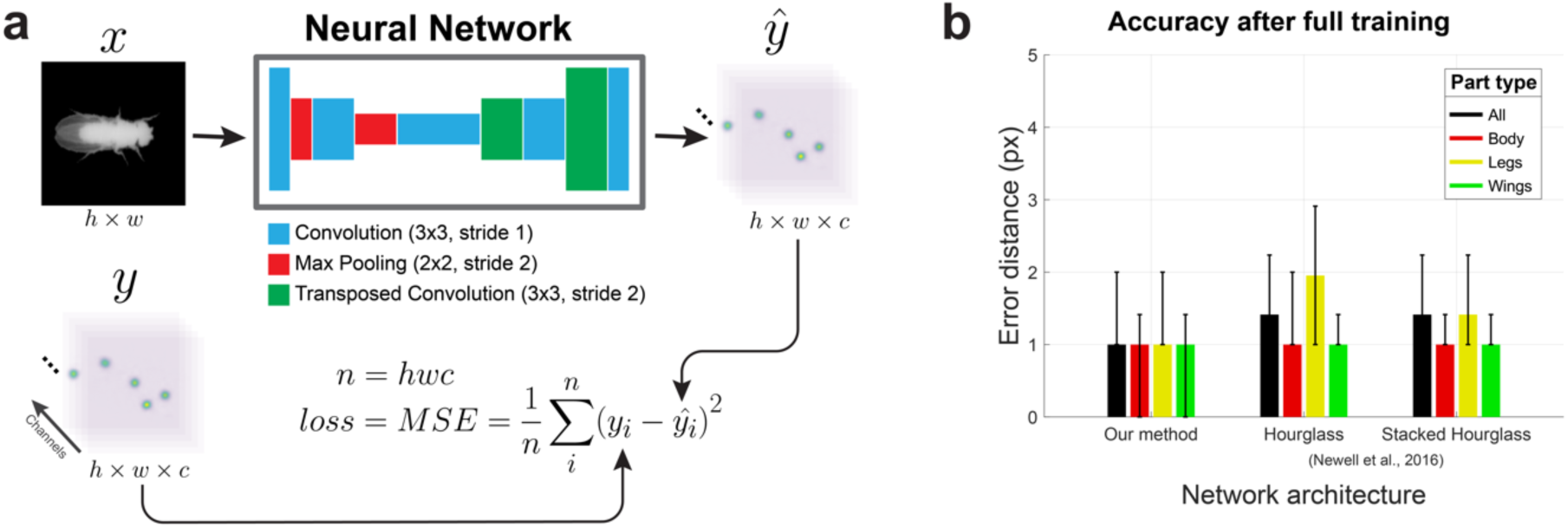
Neural network architecture comparison. (a) Diagram of our neural network architecture. Raw images are provided as input into the network, which then computes a set of confidence maps of the same height and width as the input image (top row). The network consists of a set of convolutions, max pooling and transposed convolutions whose weights are learned during training (top middle). Estimated confidence maps are compared to ground truth maps generated from user labels using a mean squared error loss function, which is then minimized during training (bottom row). (b) Accuracy comparison between architectures. We compared the accuracy of our architecture to the hourglass and stacked hourglass versions of the network described in^1^. The accuracy of our network is equivalent or better than those achieved when training with these reference architectures.

**Supplementary Figure 6:**
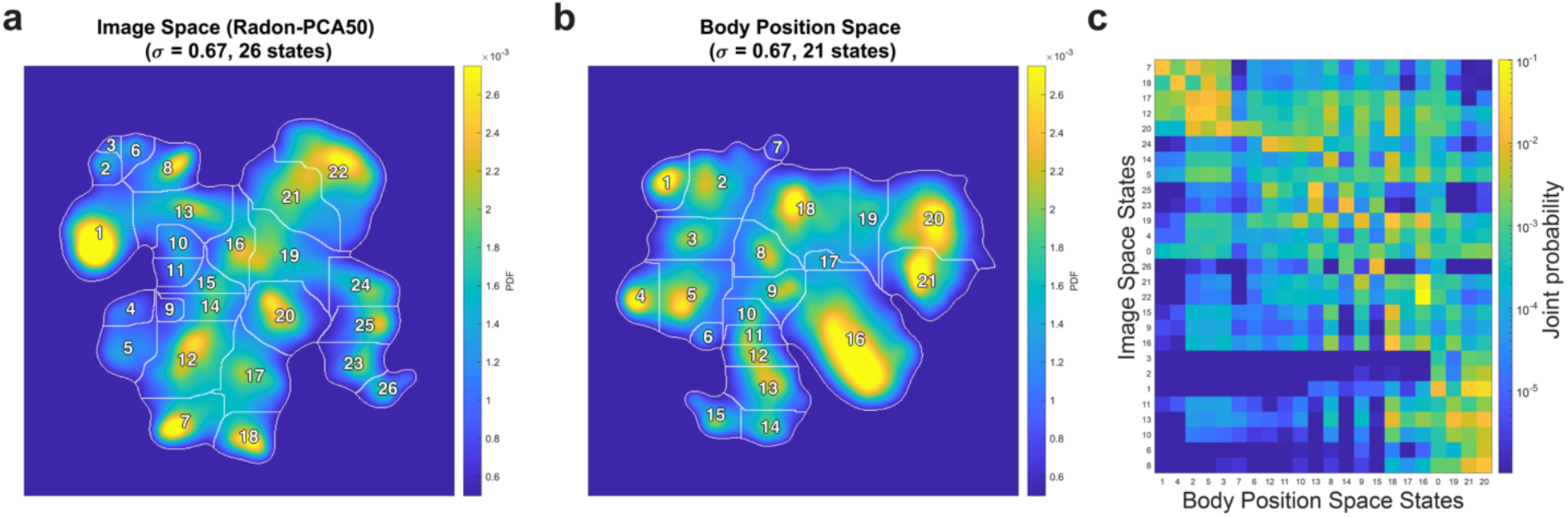
Comparison of behavioral space distributions generated from compressed images versus body part positions. (a) Behavioral space distribution from 59 male flies calculated using the original MotionMapper pipeline (data and pipeline from ^2^), including Radon-transform compression and PCA-based projection onto the first 50 principal components followed by a nonlinear embedding of the resultant spectrograms. (b) Behavioral space distribution from 59 male flies (data and pipeline from ^2^) calculated using spectrograms generated from tracked body part positions rather than PCA modes (see **Online Methods**). We note that this distribution has fewer peaks than that from (a) and a more symmetric topology (e.g in the top-left clusters, **Fig. 4c-g**). (c) Joint probability distribution of the cluster labels from (a) and (b); sorted by row and column peaks. Many clusters identified using the pixel-based representation (rows) match up with those of the position-based representation (columns), but some are distributed into newly separated clusters.

**Supplementary Movie 1:**
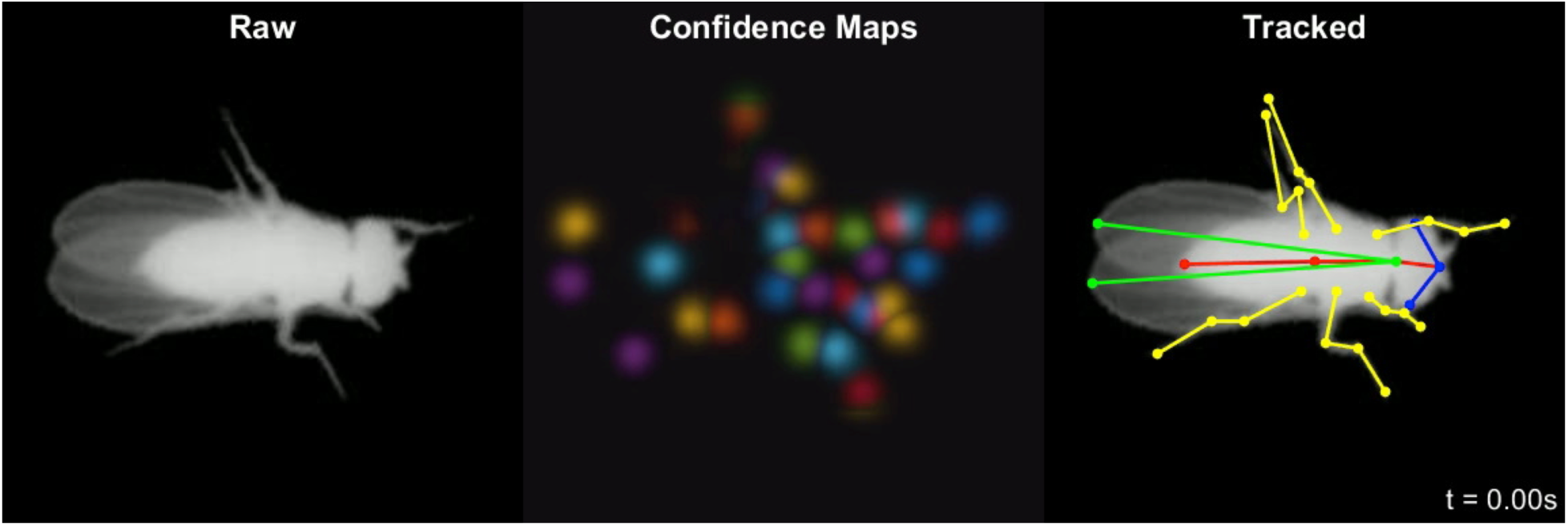
Body part tracking is reliable over long periods without temporal constraints. Raw images (left), max projection of all confidence maps (center), and tracked images (right) during a 20 second bout of free movement. Video playback at 0.2x realtime speed.

**Supplementary Movie 2:**
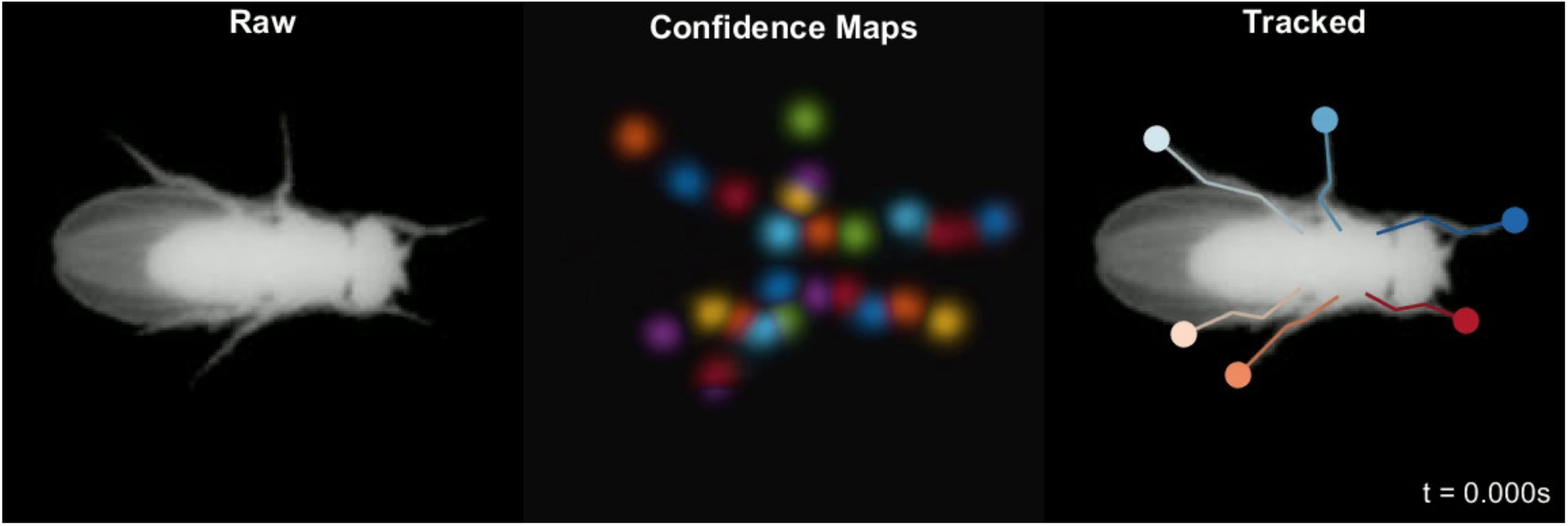
Body part tracking during freely moving locomotion. Raw images (left), max projection of all confidence maps (center), and tracked images (right) during a bout of locomotion. Video playback at 0.15x realtime speed. Video corresponds to Fig. 1d.

**Supplementary Movie 3:**
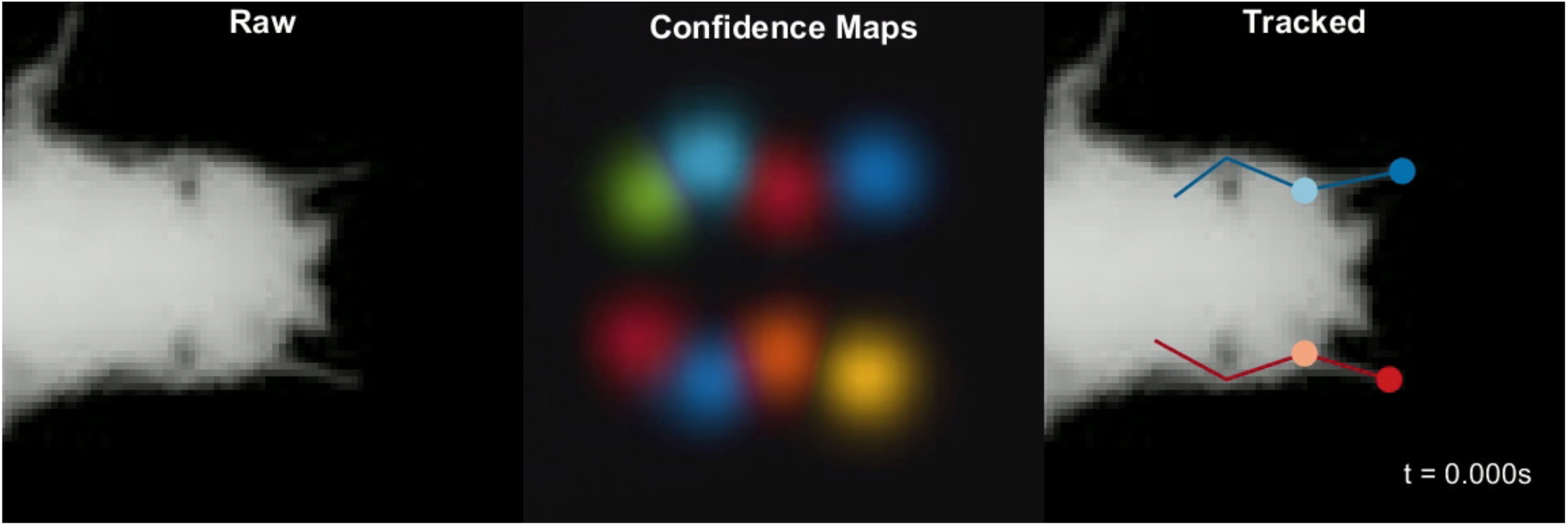
Body part tracking during head grooming. Raw images (left), max projection of all confidence maps (center), and tracked images (right) during a bout of head grooming. Video playback at 0.15x realtime speed. Video corresponds to Fig. 1e.

**Supplementary Movie 4:**
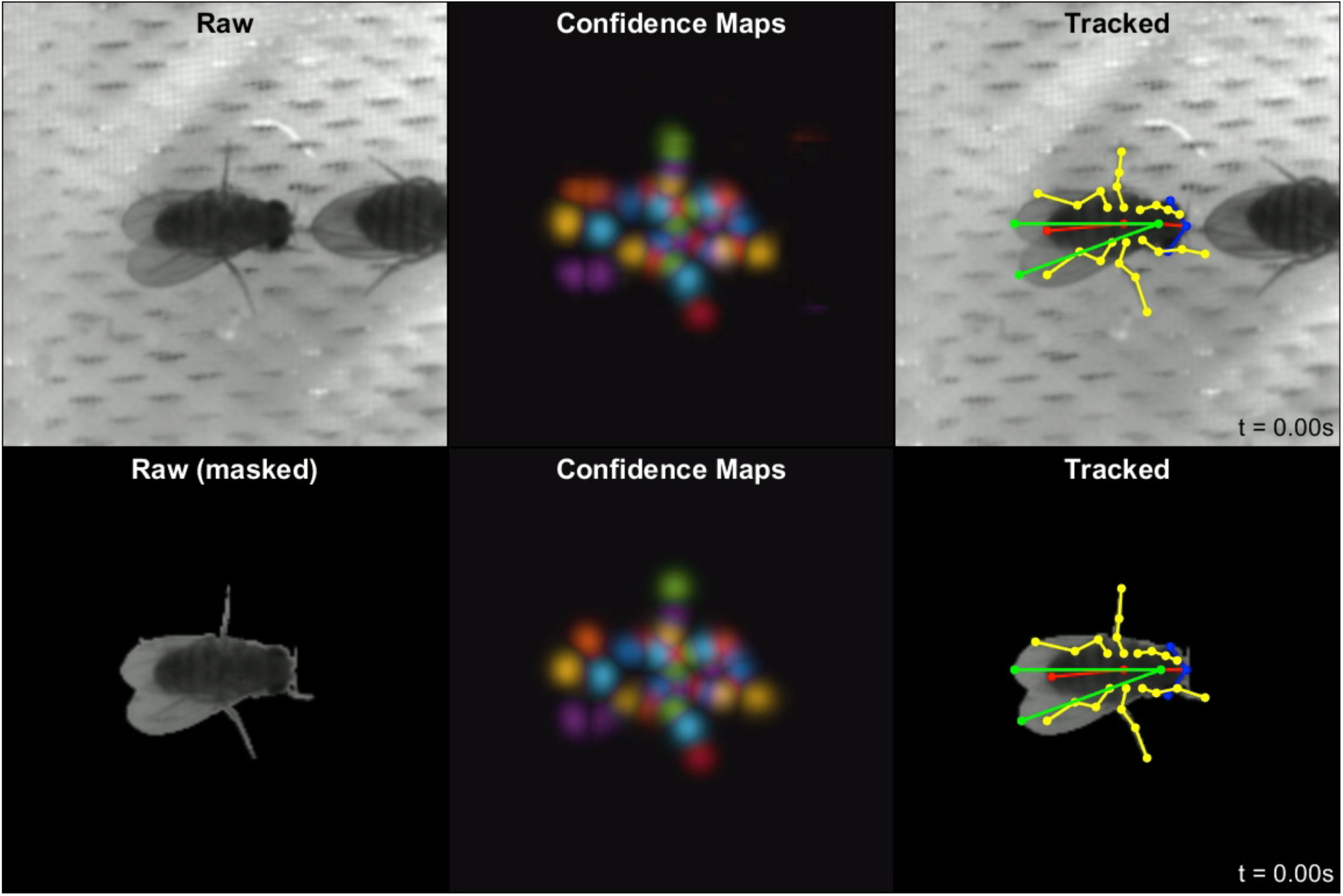
Tracking joints robustly in images with heterogeneous background and noisy segmentation. Raw images (left), max projection of all confidence maps (center), and tracked images (right) of a freely moving courting male fly. Rows correspond to results from a network trained on unmasked and masked images, respectively. Video playback at 0.2x realtime speed.

**Supplementary Movie 5:**
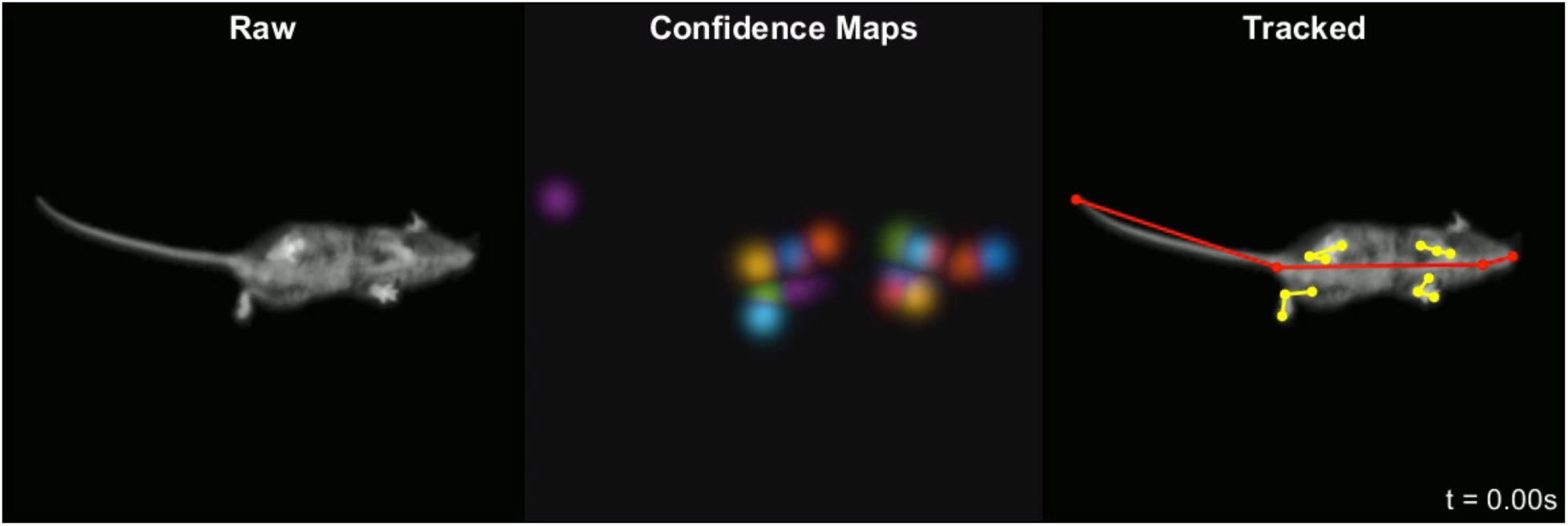
Tracking joints in freely moving rodents. Raw images (left), max projection of all confidence maps (center), and tracked images (right) of a freely moving mouse in an open field arena imaged from below through a clear acrylic floor. Video playback at 0.2x realtime speed. Tracking is reliable over time but degenerate when certain parts are occluded, such as when the animal rears.

## References

1. Anderson, D. J. & Perona, P. Toward a science of computational ethology. Neuron 84, 18–31 (2014).

2. Branson, K., Robie, A. A., Bender, J., Perona, P. & Dickinson, M. H. High-throughput ethomics in large groups of Drosophila. Nat. Methods 6, 451–457 (2009).

3. Swierczek, N. A., Giles, A. C., Rankin, C. H. & Kerr, R. A. High-throughput behavioral analysis in C. elegans. Nat. Methods 8, 592–598 (2011).

4. Deng, Y., Coen, P., Sun, M. & Shaevitz, J. W. Efficient multiple object tracking using mutually repulsive active membranes. PLoS One 8, e65769 (2013).

5. Dankert, H., Wang, L., Hoopfer, E. D., Anderson, D. J. & Perona, P. Automated monitoring and analysis of social behavior in Drosophila. Nat. Methods 6, 297–303 (2009).

6. Kabra, M., Robie, A. A., Rivera-Alba, M., Branson, S. & Branson, K. JAABA: interactive machine learning for automatic annotation of animal behavior. Nat. Methods 10, 64–67 (2013).

7. Arthur, B. J., Sunayama-Morita, T., Coen, P., Murthy, M. & Stern, D. L. Multi-channel acoustic recording and automated analysis of Drosophila courtship songs. BMC Biol. 11, 11 (2013).

8. Anderson, S. E., Dave, A. S. & Margoliash, D. Template-based automatic recognition of birdsong syllables from continuous recordings. J. Acoust. Soc. Am. 100, 1209–1219 (1996).

9. Tachibana, R. O., Oosugi, N. & Okanoya, K. Semi-automatic classification of birdsong elements using a linear support vector machine. PLoS One 9, e92584 (2014).

10. Berman, G. J., Choi, D. M., Bialek, W. & Shaevitz, J. W. Mapping the stereotyped behaviour of freely moving fruit flies. J. R. Soc. Interface 11, (2014).

11. Wiltschko, A. B. et al. Mapping Sub-Second Structure in Mouse Behavior. Neuron 88, 1121–1135 (2015).

12. Berman, G. J., Bialek, W. & Shaevitz, J. W. Predictability and hierarchy in Drosophila behavior. Proc. Natl. Acad. Sci. U. S. A. 113, 11943–11948 (2016).

13. Klibaite, U., Berman, G. J., Cande, J., Stern, D. L. & Shaevitz, J. W. An unsupervised method for quantifying the behavior of paired animals. Phys. Biol. 14, 015006 (2017).

14. Wang, Q. et al. The PSI–U1 snRNP interaction regulates male mating behavior in Drosophila. Proc. Natl. Acad. Sci. U. S. A. 113, 5269–5274 (2016).

15. Vogelstein, J. T. et al. Discovery of brainwide neural-behavioral maps via multiscale unsupervised structure learning. Science 344, 386–392 (2014).

16. Cande, J. et al. Optogenetic dissection of descending behavioral control in Drosophila. bioRxiv doi: 10.1101/230128 230128 (2018).

17. Kain, J. et al. Leg-tracking and automated behavioural classification in Drosophila. Nat. Commun. 4, 1910 (2013).

18. Machado, A. S., Darmohray, D. M., Fayad, J., Marques, H. G. & Carey, M. R. A quantitative framework for whole-body coordination reveals specific deficits in freely walking ataxic mice. Elife 4, (2015).

19. Nashaat, M. A. et al. Pixying Behavior: A Versatile Real-Time and Post Hoc Automated Optical Tracking Method for Freely Moving and Head Fixed Animals. eNeuro 4, (2017).

20. Nanjappa, A. et al. Mouse Pose Estimation From Depth Images. arXiv:1511.07611 (2015).

21. Nakamura, A. et al. Low-cost three-dimensional gait analysis system for mice with an infrared depth sensor. Neurosci. Res. 100, 55–62 (2015).

22. Wang, Z., Mirbozorgi, S. A. & Ghovanloo, M. An automated behavior analysis system for freely moving rodents using depth image. Med. Biol. Eng. Comput. (2018).:

23. Mendes, C. S. et al. Quantification of gait parameters in freely walking rodents. BMC Biol. 13, 50 (2015).

24. Mendes, C. S., Bartos, I., Akay, T., Márka, S. & Mann, R. S. Quantification of gait parameters in freely walking wild type and sensory deprived Drosophila melanogaster. Elife 2, e00231 (2013).

25. Petrou, G. & Webb, B. Detailed tracking of body and leg movements of a freely walking female cricket during phonotaxis. J. Neurosci. Methods 203, 56–68 (2012).

26. Uhlmann, V., Ramdya, P., Delgado-Gonzalo, R., Benton, R. & Unser, M. FlyLimbTracker: An active contour based approach for leg segment tracking in unmarked, freely behaving Drosophila. PLoS One 12, e0173433 (2017).

27. Toshev, A. & Szegedy, C. DeepPose: Human Pose Estimation via Deep Neural Networks. arXiv:1312.4659 (2013).

28. Tompson, J. J., Jain, A., LeCun, Y. & Bregler, C. Joint Training of a Convolutional Network and a Graphical Model for Human Pose Estimation. in Advances in Neural Information Processing Systems 27 (eds. Ghahramani, Z., Welling, M., Cortes, C., Lawrence, N. D. & Weinberger, K. Q.) 1799–1807 (Curran Associates, Inc., 2014).

29. Carreira, J., Agrawal, P., Fragkiadaki, K. & Malik, J. Human Pose Estimation with Iterative Error Feedback. arXiv:1507.06550 (2015).

30. Wei, S.-E., Ramakrishna, V., Kanade, T. & Sheikh, Y. Convolutional Pose Machines. arXiv:1602.00134 (2016).

31. Bulat, A. & Tzimiropoulos, G. Human pose estimation via Convolutional Part Heatmap Regression. arXiv:1609.01743 (2016).

32. Cao, Z., Simon, T., Wei, S.-E. & Sheikh, Y. Realtime Multi-Person 2D Pose Estimation using Part Affinity Fields. arXiv:1611.08050 (2016).

33. Tome, D., Russell, C. & Agapito, L. Lifting from the Deep: Convolutional 3D Pose Estimation from a Single Image. arXiv:1701.00295 (2017).

34. Long, J., Shelhamer, E. & Darrell, T. Fully convolutional networks for semantic segmentation. in Proceedings of the IEEE conference on computer vision and pattern recognition 3431–3440 (http://www.cv-foundation.org, 2015).

35. Ronneberger, O., Fischer, P. & Brox, T. U-Net: Convolutional Networks for Biomedical Image Segmentation. in Medical Image Computing and Computer-Assisted Intervention – MICCAI 2015 234–241 (Springer International Publishing, 2015).

36. Mathis, A. et al. Markerless tracking of user-defined features with deep learning. arXiv:1804.03142 (2018).

37. Taubin, G. Estimation of Planar Curves, Surfaces, and Nonplanar Space Curves Defined by Implicit Equations with Applications to Edge and Range Image Segmentation. IEEE Trans. Pattern Anal. Mach. Intell. 13, 1115–1138 (1991).

38. Fitzgibbon, A., Pilu, M. & Fisher, R. B. Direct Least Square Fitting of Ellipses. IEEE Trans. Pattern Anal. Mach. Intell. 21, 476–480 (1999).

39. Pérez-Escudero, A., Vicente-Page, J., Hinz, R. C., Arganda, S. & de Polavieja, G. G. idTracker: tracking individuals in a group by automatic identification of unmarked animals. Nat. Methods 11, 743–748 (2014).

40. Romero-Ferrero, F., Bergomi, M. G., Hinz, R., Heras, F. J. H. & de Polavieja, G. G. idtracker.ai: Tracking all individuals in large collectives of unmarked animals. arXiv:1803.04351 (2018).

41. Newell, A., Yang, K. & Deng, J. Stacked Hourglass Networks for Human Pose Estimation. arXiv:1603.06937 (2016).

42. Isakov, A. et al. Recovery of locomotion after injury in Drosophila melanogaster depends on proprioception. J. Exp. Biol. 219, 1760–1771 (2016).

43. Wosnitza, A., BockeMühl, T., Dübbert, M., Scholz, H. & Büschges, A. Inter-leg coordination in the control of walking speed in Drosophila. J. Exp. Biol. 216, 480–491 (2013).

44. Qiao, B., Li, C., Allen, V. W., Shirasu-Hiza, M. & Syed, S. Automated analysis of long-term grooming behavior in using a -nearest neighbors classifier. Elife 7, (2018).

45. Dombeck, D. A., Khabbaz, A. N., Collman, F., Adelman, T. L. & Tank, D. W. Imaging large-scale neural activity with cellular resolution in awake, mobile mice. Neuron 56, 43–57 (2007).

46. Seelig, J. D. & Jayaraman, V. Neural dynamics for landmark orientation and angular path integration. Nature 521, 186–191 (2015).

## References

1. Berman, G. J., Choi, D. M., Bialek, W. & Shaevitz, J. W. Mapping the stereotyped behaviour of freely moving fruit flies. J. R. Soc. Interface 11, (2014).

2. Berman, G. J., Bialek, W. & Shaevitz, J. W. Predictability and hierarchy in Drosophila behavior. Proc. Natl. Acad. Sci. U. S. A. 113, 11943–11948 (2016).

3. Chyb, S. & Gompel, N. Atlas of Drosophila Morphology: Wild-type and Classical Mutants. (Academic Press, 2013).

4. Klibaite, U., Berman, G. J., Cande, J., Stern, D. L. & Shaevitz, J. W. An unsupervised method for quantifying the behavior of paired animals. Phys. Biol. 14, 015006 (2017).

5. Tompson, J. J., Jain, A., LeCun, Y. & Bregler, C. Joint Training of a Convolutional Network and a Graphical Model for Human Pose Estimation. In Advances in Neural Information Processing Systems 27 (eds. Ghahramani, Z., Welling, M., Cortes, C., Lawrence, N. D. & Weinberger, K. Q.) 1799–1807 (Curran Associates, Inc., 2014).

6. Wei, S.-E., Ramakrishna, V., Kanade, T. & Sheikh, Y. Convolutional Pose Machines. arXiv: 1602.00134 (2016).

7. Newell, A., Yang, K. & Deng, J. Stacked Hourglass Networks for Human Pose Estimation. arXiv: 1603.06937 (2016).

8. Bulat, A. & Tzimiropoulos, G. Human pose estimation via Convolutional Part Heatmap Regression. arXiv: 1609.01743 (2016).

9. Cao, Z., Simon, T., Wei, S.-E. & Sheikh, Y. Realtime Multi-Person 2D Pose Estimation using Part Affinity Fields. arXiv: 1611.08050 (2016).

10. Kingma, D. P. & Ba, J. Adam: A Method for Stochastic Optimization. arXiv: 1412.6980 (2014).

11. Morel, P. Gramm: grammar of graphics plotting in Matlab. JOSS 3, 568 (2018).

12. Mendes, C. S., Bartos, I., Akay, T., Márka, S. & Mann, R. S. Quantification of gait parameters in freely walking wild type and sensory deprived Drosophila melanogaster. Elife 2, e00231 (2013).

13. Isakov, A. et al. Recovery of locomotion after injury in Drosophila melanogaster depends on proprioception. J. Exp. Biol. 219, 1760–1771 (2016).

14. Baum, L. E., Petrie, T., Soules, G. & Weiss, N. A Maximization Technique Occurring in the Statistical Analysis of Probabilistic Functions of Markov Chains. Ann. Math. Stat. 41, 164–171 (1970).

15. Viterbi, A. Error bounds for convolutional codes and an asymptotically optimum decoding algorithm. IEEE Trans. Inf. Theory 13, 260–269 (1967).

16. Maaten, L. van der & Hinton, G. Visualizing Data using t-SNE. J. Mach. Learn. Res. 9, 2579–2605 (2008).

## References

Newell, A., Yang, K. & Deng, J. Stacked Hourglass Networks for Human Pose Estimation. In Computer Vision – ECCV 2016 483–499 (Springer International Publishing, 2016).

Berman, G. J., Choi, D. M., Bialek, W. & Shaevitz, J. W. Mapping the stereotyped behaviour of freely moving fruit flies. J. R. Soc. Interface 11, (2014).

